# Meta-analysis reveals the tempo of evolutionary parallelism of local adaptation between native and introduced ranges of invasive plant species

**DOI:** 10.1101/2025.09.04.674372

**Authors:** Romane Normand, Alexis Heckley, Kathryn Hodgins, Samantha Grover, Tim Connallon, Akane Uesugi

## Abstract

Invasive species are valuable systems for evaluating evolutionary predictability, as populations in native and introduced ranges evolve separately, yet often encounter similar environmental challenges that select for parallel patterns of local adaptation across each range. However, it remains unclear how pervasive and strong such parallelism is and how rapidly it evolves. To address these questions, we first extended cline theory to predict evolutionary patterns of trait cline parallelism between ranges. We then carried out a meta-analysis of clinal divergence in native and introduced populations of the same plant species and evaluated the tempo of trait cline parallelism between ranges. We found that clines in introduced ranges were somewhat shallower than native range clines, and directions of clinal divergence were, on average, partially aligned between ranges, with extensive variation in parallelism among traits and species, including counter-clines between native and introduced ranges. The degree of evolutionary parallelism of clinal divergence strongly increased, and the prevalence of counter-clines decreased, with the time since introduction, where greater parallelism in older introductions was primarily caused by increased alignment in the direction of clinal divergence between ranges rather than changes in their relative magnitudes of divergence. We argue that these observations are consistent with a two-phased process of cline evolution across introduced ranges. In the first phase, misaligned clines between introduced and native ranges can arise during range expansions, often owing to demographic and genetic constraints that initially hinder the evolution of local adaptation within the new range. During the second phase, the evolution of local adaptation eventually resolves early instances of maladaptation across introduced ranges, leading to strong trait cline parallelism between native and introduced ranges.

## Introduction

A general understanding of the process of adaptation must address several broad questions about how ecology, evolution and genetics interact in natural populations. How predictable is evolution? Do shared environmental conditions lead to reliably parallel evolutionary outcomes, or are there multiple genetic or phenotypic solutions to the same environmental challenges [1,2]? Do populations typically adapt rapidly over ecological timescales [3,4], or do they experience sustained evolutionary lags in adaptation to their environments, owing to limited genetic variation or other constraints [5,6]? What are the key environmental variables—abiotic and biotic—that determine natural selection and the direction of evolutionary change [7,8]? Each of these questions is difficult to answer, especially in natural systems.

Introduced species provide exceptional study systems for addressing questions about adaptation, including its predictability, rate, and genetic basis [9,10]. In particular, shared environmental factors between the native and introduced ranges of a species can favor the evolution of parallel patterns of clinal divergence between ranges (“parallelism”), in which traits or the frequencies of genetic variants show similar geographical patterns of divergence along spatial gradients in temperature, aridity, or other environmental variables (e.g., [11–13]). In other cases, evolutionary contingency or environmental differences between the ranges can select for different patterns of clinal divergence between native versus introduced populations (e.g., [14,15]). Non-adaptive mechanisms of cline formation can likewise yield different patterns of spatial divergence across different ranges of a species [16,17]. In addition, the age of an introduction constrains the time available for adaptation to occur in the new range. In some cases, rapid evolutionary diversification across the new range may keep pace with the range expansion itself, leading to substantial local adaptation within decades of the introduction [18,19]. In other cases, introduced populations may experience evolutionary lags, with protracted periods of suboptimal local adaptation across their introduced ranges [20,21]. Finally, these phenomena may play out differently among trait types [22], which may show differences in the strength or direction of selection that they experience, or in their genetic capacities for evolutionary change.

Comparisons of native and introduced populations of the same species offer two major advantages for evaluating the evolutionary rate and predictability of local adaptation, as implied by parallel patterns of clinal divergence in each range [16,17]. First, such studies have a natural, paired design, akin to case-control experiments, in which the evolution of a common set of traits can be directly compared across the native and introduced ranges [23]. Such comparisons are particularly powerful when carried out using individuals reared in common garden environments [24], which allow evolved differences among populations to be isolated from environmental factors. One can then directly contrast the direction and steepness of geographic clines across the introduced range, whose populations are relatively recent and potentially maladapted to their local conditions, with the native range, whose populations have had extensive time to evolve and are presumably better adapted to local conditions [25,26]. Second, introduction dates can often be approximated using historical records, which provide opportunities to quantitatively test whether patterns of parallelism between ranges increase with the amount of time since the introduction [21,27]. If selection favors similar trait–environment relationships in both ranges, older introductions should exhibit greater parallelism than more recent introductions [28].

Studies of clinal divergence in native versus introduced populations of the same species have progressed to a point where we now have many high-quality case studies spanning a diverse range of plant taxa [22]. These studies, if analyzed in a common framework, provide an opportunity to test hypotheses about local adaptation, including the degree and tempo with which parallelism evolves between native and introduced portions of species’ ranges. With this goal in mind, we conducted a systematic search for common-garden trait cline studies in plants and identified 35 studies, spanning 24 species and 513 traits, that report cline estimates in both the native and introduced portions of the species’ range. Our dataset expands upon earlier meta-analyses of trait clines in native versus introduced plant populations by incorporating studies published over the last 15 years ([22] includes studies published up to 2011).

We begin by presenting an extension of classical cline theory that generates quantitative predictions for the evolution of cline parallelism between native and introduced ranges of a species. We then carry out an analysis of published cline data to address three questions: (1) How predictable is local adaptation, as implied by evolutionary parallelism of trait clines between native and introduced ranges? (2) How rapid is the evolution of local adaptation, as implied by the interaction between the time since introduction and the degree of parallelism between introduced and native populations? (3) To what extent do different trait categories (e.g., morphological, phenological, and defense traits) show parallelism between the native and introduced ranges?

### Trait clines in native versus introduced ranges: Theoretical predictions

To facilitate interpretation of the meta-analysis results, we first derived theoretical predictions for trait cline parallelism between ranges for a set of quantitative traits contributing to local adaptation. The main results of our analysis, which build upon classic cline models from García-Ramos and Kirkpatrick [29], are summarized below and extended results are provided in the Supporting Information (SI). Following García-Ramos and Kirkpatrick [29], we make several assumptions: (1) the optimum of each trait changes linearly across continuous geographic gradients (e.g., latitude) in the native and introduced ranges of a species; (2) conditions in the native range are sufficiently stable that native-range populations are locally adapted and at equilibrium between local selection and gene flow; and (3) introduced range populations are initially displaced from equilibrium but, over time, eventually evolve to migration-selection balance across the new range.

For simplicity, we compartmentalize the evolution of clines in the introduced range into two phases. *Phase one* coincides with the range expansion itself, which involves the establishment of one or more founder populations, followed by geographic expansion of the species from these founder sites and across the new range. Prior work has shown that clines can emerge by several processes during this phase. Adaptive clines can emerge due to strong selection and the evolution of rapid local adaptation during a geographic expansion (e.g., [30,31]) or by preferential establishment of populations that are pre-adapted to the local conditions at different points across the new range [32,33]. Nonadaptive clines can arise by chance events during the range expansion, including population structure arising from multiple, random introductions at different locations with the new range followed by incomplete mixing upon contact between them [34], and by strong genetic drift caused by serial bottlenecks during spatial expansions across the introduced range [22]. These factors can collectively lead to the presence of many well-defined clines near the onset of a species’ introduction, and we treat such clines as initial conditions in our models.

During *phase two*, which commences once introduced populations have reached large and relatively stable sizes, we model the evolutionary dynamics of clines as a deterministic process shaped by migration and local selection across the new range. Importantly, the initial conditions of phase two should depend on the specific mechanism of cline formation during phase one. In cases where the initial clines are adaptive, local selection during phase two will simply reinforce and potentially strengthen the clines that arose during phase one. However, in cases where the initial clines are maladaptive (e.g., a potentially substantial subset of clines arising by drift or population structure caused by complex introductions), evolution of local adaptation during phase two is expected to eliminate or reverse the direction of the initial clines that arose during phase one.

Let *t* represent the number of generations since completion of the range expansion (*t* = 0 represents the onset of phase two). For a single trait, let *B_N_* and *B_I_* represent the rates of change (slopes) of the trait optima across the native and introduced ranges, respectively (see the dashed lines in Fig. 1A), whereas *b_N_* and *b_I_* represent cline slopes of the trait (solid curves in Fig. 1A). Following García-Ramos and Kirkpatrick [29], we assume the trait is normally distributed with unit variance at each location in a range, and the trait heritability is constant across each range. Local selection is a Gaussian function of the trait value, with the optimum changing across each range and the strength of stabilizing selection remaining constant. With strong local selection relative to gene flow, equilibrium trait clines closely track the optimum. The native range cline is, thus, approximately *b_N_* = *B_N_* (see Fig. 1A), while the introduced range cline is approximately:

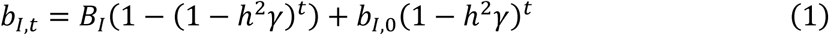

**Figure 1.**
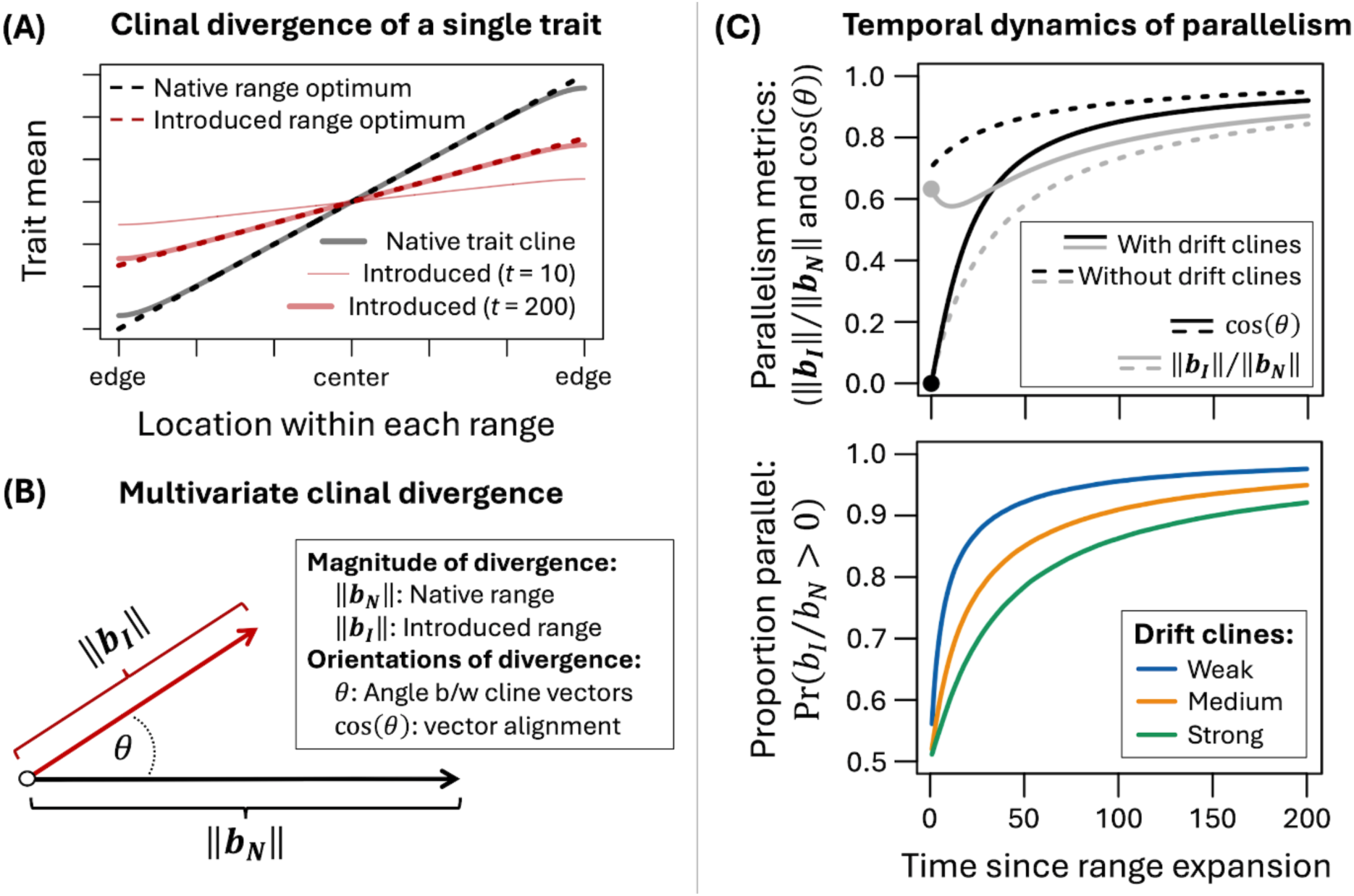
Theoretical predictions for trait cline parallelism between ranges. **Panel A:** Examples of single trait clines in a native (black) and introduced range (red). The native range trait cline is evaluated at equilibrium, while the introduced trait cline is evaluated after *t* = 10 and *t* = 200 generations following the range expansion with no initial cline in the introduced range. Selection for local adaptation is assumed to be strong relative to the swamping effects of gene flow, leading to a close match between trait clines and optima at equilibrium (see SI Appendix 1). Extensions of the model and further results are presented in SI Appendix 1-2. **Panel B:** Metrics of multivariate trait clines across the native and introduced ranges of a species. The lengths of black and red arrows show the magnitudes of clinal divergence in the native and introduced ranges, respectively. The cosine of the angle between cline slope vectors (−1 ≤ cos(*θ*) ≤ 1) measures the alignment of cline orientations between ranges (see the text for elaboration). **Panel C:** Evolution of increased parallelism between native and introduced ranges that span similar environmental conditions. The top panel shows the tempo of convergence between ranges for the magnitudes and orientations of native and introduced trait cline vectors (‖***b_I_***‖⁄‖***b_N_***‖ and cos(*θ*) approach one over time). The bottom panel shows the proportion of traits exhibiting parallel clines between ranges (the proportion of traits with *b_I_*⁄*b_N_* > 0, as opposed to counter-clines, where *b_I_*⁄*b_N_* < 0). aative range clines are at migration-selection equilibrium, while introduced clines gradually evolve towards equilibrium following the range expansion. Total clinal divergence is determined by many genetically independent traits (*n* = 10^5^), with the product of heritability and stabilizing selection for each trait (ℎ^2^*γ*) drawn from an exponential distribution with mean of 0.02. Cline optimum slopes for the traits follow a standard normal distribution, assumed here to be identical between ranges, which leads to perfect parallelism at equilibrium (‖***b_I_***‖⁄‖***b_N_***‖ = cos(*θ*) = 1 with large *t*). Genetic drift during the range expansion (see [22]) generates random initial cline slopes and initially poor alignment between the native and introduced ranges. The top panel compares the dynamics of parallelism with strong drift-induced initial trait clines (solid curves, where var(*b_I_*_,0_) = 0.4 var(*B_N_*)) versus parallelism under selection and migration with no initial clines (broken curves, where *b_I_*_,0_ = 0). The grey and black circles show the predicted initial states with drift clines: 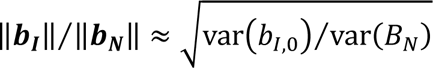 and cos(*θ*) ≈ 0 (SI Appendix 5). The bottom panel shows three degrees of drift-induced clines: weak drift clines (var(*b_I_*_,0_) = 0.01 var(*B_I_*)), medium drift clines (var(*b_I_*_,0_) = 0.1 var(*B_I_*)), and strong drift clines (var(*b_I_*_,0_) = 0.5 var(*B_I_*))).

(SI Appendix 1), where ℎ^2^ is the trait’s heritability, *γ* is the strength of stabilizing selection (ℎ^2^ and *γ* are subject to the constraints: 0 ≤ ℎ^2^ ≤ 1; 0 < *γ* < 1; see SI Appendix 1), and *b_I_*_,0_ is the initial trait cline slope immediately following the range expansion (the slope at time *t* = 0). The ratio of trait cline slopes over time is therefore *b_I_*_,*t*_⁄*b_N_* ≈ *b_I_*_,*t*_⁄*B_N_*. If the initial cline in the introduced range is small relative to the native range cline (i.e., |*B_N_*| ≫ |*b_I_*_,0_|), then the ratio of introduced to native range trait cline slopes will be nearly zero shortly after the range expansion (e.g., *b_I_*_,*t*_⁄*b_N_* ≈ 0 when *t* is small; see Fig. 1A) and then converge over time to *b_I_*⁄*b_N_* = *B_I_*⁄*B_N_*, which ultimately reflects the similarity of local selection between the ranges.

***Perfect parallelism*** of trait clines is expected at equilibrium if spatial changes of the optima are identical between ranges (at equilibrium, *b_I_*⁄*b_N_* = 1 when *B_I_* = *B_N_*) and all else is equal. ***Partial parallelism*** is expected when geographic shifts in the trait optima occur in the same direction in each range but differ in magnitude (i.e., *b_I_*⁄*b_N_* > 0 and *b_I_*⁄*b_N_* ≠ 1, at equilibrium, if *B_I_*⁄*B_N_* > 0 and *B_I_*⁄*B_N_* ≠ 1), or when there are differences between ranges in trait heritability or the relative strengths of selection and gene flow, which can lead to persistently unequal cline slopes between ranges despite similar environmental conditions (see SI Appendix 2). ***Antiparallelism*** (i.e., “counter-clines”, where *b_I_*_,*t*_⁄*b_N_* < 0) can occur in non-equilibrium populations when clines arise by drift or population structure during the range expansion (see [22]) or from indirect responses to selection on genetically correlated traits (see SI Appendix 6; for similar concepts but somewhat different terminology, see [35]). However, at equilibrium, counter-clines are only expected when trait optima shift in opposite directions between the ranges (*B_I_* and *B_N_* have opposite signs). Overall, a strong empirical pattern of parallelism is eventually expected (i.e., in sufficiently old introductions) when patterns of spatially varying selection are similar between native and introduced ranges. Deviations from perfect parallelism can arise from evolutionary lags to the equilibrium in the introduced range, divergent patterns of selection between ranges, or systematic differences between ranges in genetic or demographic constraints to local adaptation (SI Appendix 2; also see [22,29,36]).

Organisms are, of course, comprised of many traits that collectively respond to selection, leading to multivariate clinal divergence. Multivariate parallelism between ranges of a species can be quantified by comparing vectors of trait cline slopes from each range (Fig. 1B). Letting ***b_I_*** = (*b_I_*_,1_, *b_I_*_,2_, …, *b_I_*_,*n*_) and ***b_N_*** = (*b_N_*_,1_, *b_N_*_,2_, …, *b_N_*_,*n*_) represent the introduced- and native-range vectors of cline slopes for a common set of *n* traits, the total amount of clinal divergence in each range is captured by the Euclidean norms of the vectors 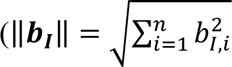 and 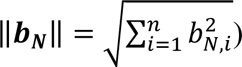 and alignment of the multivariate orientations of clinal divergence in each range, as captured by the cosine of the angle between the cline vectors (their “cosine similarity”):

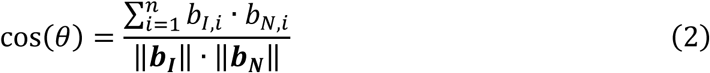

(see Fig. 1B), where cos(*θ*) = 1 indicates perfect alignment of cline orientations, 0 < cos(*θ*) < 1 indicates partial alignment, and cos(*θ*) < 0 indicates misalignment between the ranges. Perfect parallelism between ranges occurs when ‖***b_I_***‖⁄‖***b_N_***‖ = cos(*θ*) = 1, and deviations from perfect parallelism can occur when ranges differ in their magnitudes and/or orientations of clinal divergence (‖***b_N_***‖ ≠ ‖***b_I_***‖ and/or cos(*θ*) < 1).

What patterns of multivariate parallelism should we expect when native and introduced ranges span similar environmental conditions? If traits contributing to local adaptation are genetically independent of one another, local selection is strong relative to gene flow, and trait clines are initially absent following the range expansion (*b_I_*_,0_ = 0 for each trait), then the model makes three predictions (see SI Appendix 3 and the broken lines in Fig. 1C). First, none of the traits will show counter-clines between ranges (*b_I_*_,*t*_⁄*b_N_* ≥ 0 for all traits at all times). Second, the ratio of cline magnitudes shows a strong increase over time, with an initial value of ‖***b_I_***‖⁄‖***b_N_***‖ = 0 and a final value of ‖***b_I_***‖⁄‖***b_N_***‖ ≈ 1 if changes in the trait optima are the same between ranges (or more generally, 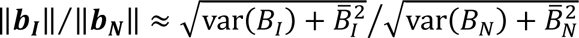 at equilibrium, with *B̄_I_*, *B̄_N_*, var(*B_I_*) and var(*B_N_*) representing the means and variances of optimum slopes of the ranges). Third, the alignment of cline vectors increases from an initial value of 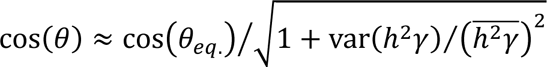 to an equilibrium value of cos(*θ*_eq._), where var(ℎ^2^*γ*) represent the mean and variance among traits for the compound parameter ℎ^2^*γ*. Variation in ℎ^2^*γ* causes heterogeneity among traits in the rate at which they respond to selection, which dampens the initial alignment of clinal divergence between the ranges relative to the alignment at equilibrium. However, this effect is usually small and the total change in cos(*θ*) over time is, therefore, small relative to the total change in ‖***b_I_***‖⁄‖***b_N_***‖. Similar patterns are expected when the process of local adaptation begins during the range expansion, in which case the initial trait clines should be proportional to the initial rates of change in cline slopes due to selection (∼ℎ^2^*γB_I_* for a single trait in our model).

These three predictions can change substantially when maladaptive counter-clines arise during the range expansion, either by chance or due to genetically correlated evolutionary responses to selection on other traits. For example, if clines initially evolve under pure genetic drift during the range expansion (as in [22]), then their initial slopes may reasonably be approximated as normally distributed with mean of zero and variance (denoted var(*b_I_*_,0_)) determined by the demography of the range expansion and the heritability of the trait (see SI Appendix 4). While we do not explicitly model the formation of clines by drift, the phenomenon is easily incorporated into our model by exploring how different values of var(*b_I_*_,0_) affect the temporal patterns of multivariate parallelism between ranges (SI Appendix 5). Returning to our model of genetically independent clines (as above), we find that drift-induced clines, firstly, generate a high proportion of counter-clines between ranges involving recent introductions (∼50% of traits, initially; Fig. 1C, bottom panel). This proportion then declines over time as introduced populations locally adapt to environmental conditions that parallel those in the native range. Second, emergence of clines during the range expansion can lead to substantial clinal divergence in very recent introductions (e.g., the solid grey curve in Fig. 1C, top panel, where the initial ratio of magnitudes caused by drift-induced clines is 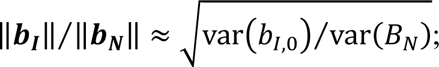; see SI Appendix 5). In such cases, the ratio of cline magnitudes may show relatively modest change over time. Third, the alignment between native and introduced trait cline orientations shows more pronounced change over time. When initial clines are drift-induced, we see a shift from initially no alignment (cos(*θ*) = 0) to strong alignment at equilibrium (e.g., cos(*θ*) = 1 at equilibrium when environmental conditions are identical between the ranges, the solid black curve in Fig. 1C, top panel), where the total change in cos(*θ*) over time can be much greater than the change in ‖***b_I_***‖⁄‖***b_N_***‖. Similar patterns can occur when genetic correlations between traits lead to maladaptive counter-clines in recent introductions, which can produce (depending on the timescale) greater change in cos(*θ*) than in ‖***b_I_***‖⁄‖***b_N_***‖ (see SI Appendix 6).

In summary, temporal patterns of parallelism in clinal divergence should reflect the similarity of environmental conditions between native and introduced ranges (which determines whether selection favors the evolution of parallel clines), the role of chance and genetic constraints to adaptation within introduced ranges (which can lead to maladaptive clines, particularly so in recent introductions), and the amount of time that introduced populations have had to adapt to conditions across their new ranges. To the extent that chance and constraints affect the initial emergence of trait clines, we expect counter-clines between native and introduced ranges to be more common in recent relative to old introductions, and overall patterns of cline parallelism between ranges should shift from weak to strong over time.

## Methods

### Meta-analysis data

We compiled the meta-analysis data following PRISMA guidelines ([37]; see SI Fig. S1), and included studies reporting trait clines in both the native and introduced ranges of the study species. Eligible articles were identified from searches of Web of Science, Scopus, and PubMed (initial searches were carried out in December 2022 and repeated in May 2025). Keywords were a combination of synonyms related to clines (e.g., “clinal variation”), introduced species (e.g., “introduced”), plant (e.g., “weed”), and the term “adapt*” (see SI Fig. S1 for details). We included studies using common garden experiments published in academic journals with titles and abstracts in English. Studies cited in identified articles were added to our database if they met the inclusion criteria.

When possible, we obtained raw data published online or provided by authors. When raw data were not available, we extracted mean trait values and standard errors from figures using ImageJ [38]. Studies with less than four populations in each range were excluded from the analysis. We also excluded studies focusing on ‘expanded’ ranges, where the native range has recently expanded within the same continent [39]. A total of 35 studies met our inclusion criteria (SI Box S1). Some studies grew plants under multiple common garden treatments (e.g., different water or nutrient regimes) or implemented common garden experiments at different sites. These “experiments” were analyzed separately because trait values could not be directly compared across experiments. When multiple introduced ranges were tested against a single native range, we considered each pair of contrasts as a separate “experiment”. The final dataset includes paired (native and introduced) cline estimates for 513 traits, from 84 experiments, spanning 24 species and 10 plant families (SI Table S1). Each trait was classified into one of the following categories (SI Table S2): morpho-physiology (structural features and functional processes), reproduction (investment in offspring production), size (biomass and growth traits), phenology (timing of developmental stages) and defense (physical and chemical traits that resist herbivory).

### Estimation of the number of generations since introduction

To estimate species’ introduction times, we used the year of introduction to the study area when it was provided by authors. When that information was not available, we used the first record of introduction reported by Seebens *et al*. [40], for countries matching the source populations of each species in our dataset. When records for the source country were unavailable, we used records from the closest country on the same continent. For studies involving populations sampled from multiple countries, we used the average year of the first record across the countries. Years since introduction were calculated as the year of seed collection minus the year of first record. When seed collection dates were not available, we used the year of the common garden experiment. The number of generations since introduction was calculated as [years since introduction]/[generation time], with species’ generation times (in years) obtained from [41]. When generation time data were not available, we approximated generation time as *T_z_* = *α* + 0.5(*ω* − *α*), where *α* is the species’ age at reproductive maturity and *ω* is the typical lifespan [42], each obtained from the TRY database [43]. With accurate measures of *α* and *ω*, *T_z_* provides a good approximation of generation time [44].

### Effect size calculations

All statistical analyses were performed with the R software v 4.4.1 [45].

#### Estimation of individual trait cline slopes

For each experiment, we estimated cline slopes using linear models with the population-level trait means as the response variable and the environmental gradient the authors of the original studies focused on as the explanatory variable. These environmental gradients included latitude, principal components of BioClim data, temperature, elevation, aridity, altitude, and longitude. We also analyzed the data using a single shared environmental predictor across the set of experiments (e.g., absolute latitude, temperature, or precipitation, with the latter two predictors calculated following Atwater *et al*. [46]). Within each experiment, traits were standardized (mean = 0; standard deviation = 1) across both native and introduced populations before model fitting. Traits with highly skewed distributions were square root or log transformed prior to standardization to improve residual normality. For each trait, population means were used to obtain a native slope (*b_N_*), an introduced slope (*b_I_*), and their standard errors.

#### Multivariate parallelism

We estimated multivariate parallelism in 73 experiments with more than one trait. For each experiment, we compiled vectors of trait cline estimates for the introduced and native range (***b_I_*** and ***b_N_***), from which we calculated vector norms (‖***b_I_***‖ and ‖***b_N_***‖) and the cosine between vectors (cos(*θ*)) using eq. (2) (also see [47,48]). We analyzed log-transformed ratio of magnitudes, ln(‖***b_I_***‖/‖***b_N_***‖), where ln(‖***b_I_***‖/‖***b_N_***‖) = 0 indicates equal magnitudes between ranges.

Following Poissant et al. [49], uncertainty around the point estimates of cos(*θ*) and ln(‖***b_I_***‖/‖***b_N_***‖) was estimated by bootstrapping the original data by randomly selecting individual samples with replacement. For each bootstrapped sample, we used population means of trait values to calculate the parameters using the methods described above. In cases where only the population means and standard errors were available (but not raw data), we simulated trait means for each population by pseudo-random sampling from a normal distribution with the mean corresponding to the point estimate of the mean for the population and standard deviation corresponding to its standard error. Each set of simulated means across populations within a range was used to calculate cline slopes, from which we recalculated multivariate parameters of parallelism. We repeated this process 10,000 times to approximate confidence intervals for these parameters.

#### Univariate parallelism

Our univariate analysis focused on traits with statistically significant clines in at least one range (270 of 513 traits in the dataset), as those without a significant cline in either range are irrelevant to tests of parallelism. We used ratios of cline slopes (i.e., *b_I_*/*b_N_*) because they facilitate comparisons of effect sizes between the wide assortment of traits and study systems in the dataset, regardless of the original units of measurement of the traits or the environmental gradients chosen by the authors. Following Friedrich *et al*. [50], we calculated the relative magnitudes of univariate clinal divergence using the natural logarithm of cline slope ratios (ln(*b_I_*/*b_N_*)), which allows parametric statistical analyses of meta-analysis effect sizes. 95% confidence intervals for the mean ln(*b_I_*/*b_N_*) of each trait were calculated using equations presented by Friedrich *et al*. [50].

### Meta-analysis

#### Overview

Meta-analyses were performed using the function *rma.mv* from the package ‘metafor’ [51]. Parameters in the meta-analytical models were estimated using restricted maximum likelihood [52], with Knapp-Hartung adjustments used to calculate the confidence interval of pooled effects [53]. We performed multilevel random-effects models to account for the hierarchical structure of non-independence in our dataset (multiple traits in multiple experiments within single studies; multiple studies examine the same species). We used the function *var.comp* from the package ‘dmetar’ [54] to calculate the total heterogeneity (I^2^), and heterogeneity at each level (I^2^_level 2_ and I^2^_level 3_). We tested whether residual heterogeneity was due to sampling error only, using the QE statistic, with significant QE indicating that residual heterogeneity can be explained by factors not included in the model [51].

#### Multivariate analysis

The meta-analysis of multivariate parameters estimated pooled effect sizes and 95% confidence intervals for cos(*θ*) and ln(‖***b_I_***‖/‖***b_N_***‖), with random effects of study identity nested within a species. We tested how multivariate parameters change with time since introduction by including generations since introduction as a moderator in the meta-analysis model. We also included the following potential sources of variation as moderators: 1) the type of environmental gradients authors used to estimate clines (‘*gradient type*’ = latitude, elevation, aridity, temperature, or BioClim PC axis); 2) the proportion of the environmental gradient that is shared between native and introduced ranges (‘*gradient overlap*’), calculated as: [the amount of overlap in environmental gradient between introduced and native ranges] / [total gradient covered by both ranges]; 3) whether populations were introduced into the same hemisphere as their native counterparts or a different one (‘*hemisphere contrast*’), as transitions within the same hemisphere (e.g., North America to Europe, or South Africa to Australia) may encounter more similar biotic environments than transitions between hemispheres (e.g., due to different species assemblies associated with the Laurasian vs. Gondwanan supercontinents); 4) mating system of the species (‘*mating system*’), categorized as self-incompatible, self-incompatible with self-compatible populations reported in introduced ranges, self-compatible, apomictic and clonal (TRY database); 5) the number of traits (‘*trait number*’) examined and included in multivariate analysis; and 6) the signal-to-noise ratio (‘*S:N*’) of cline slope estimates, calculated as *S*: *N* = ‖***b_I_***‖ · ‖***b_N_***‖⁄(‖***SE_I_***‖ · ‖***SE_N_***‖), where ***b_I_*** and ***b_N_*** are vectors of cline slope estimates (as above), ***SE_I_*** and ***SE_N_*** are corresponding vectors of the standard errors of the slope estimates, and double-vertical bars denote vector norms [55]. This last variable was included because estimation error in elements of a pair of vectors tends to downwardly bias estimates of cosine similarity between them and upwardly bias estimates of multivariate magnitudes [56]. This bias is expected to be large for experiments with low signal-to-noise, such that cos(*θ*) approaches zero, on average, when estimates of clinal divergence are of similar magnitude as error in the estimates. By including *S*: *N* as a covariate in our model, we control for the possibility that associations between cos(*θ*) estimates and the time since introduction are not confounded by heterogeneity among studies in precision of cline estimates.

We used the function *glmulti* (package ‘glmulti’ [57]) to select the best model based on Akaike information criterion with small sample size correction (AICc). Models within ΔAICc < 2 were considered competitive [58], and we compared the relative importance of the terms found among the competitive models (i.e., the sum of the relative evidence weights of all models in which the term appears [48]). The final model structure was determined by choosing the competitive model with terms with high importance value.

#### Univariate analysis

We tested whether the proportion of clines going in the same direction (*b_I_*/*b_N_* > 0) changed with the number of generations since introduction (as predicted by theory; Fig. 1C) using a generalized linear model (*glm*) with generations-since-introduction as the fixed effect, a binomial error distribution and a logit link function, and a binary response variable (0 = [*b_I_*/*b_N_* <0] and 1 = [*b_I_*/*b_N_* > 0]).

The subsequent univariate meta-analysis focused on positive ratios of clines (traits where *b_I_*/*b_N_* > 0), which indicate parallelism in a true sense and because logarithm is defined in such cases (we note that analyses of the absolute value of *b_I_*/*b_N_*, using the full set of clines that include *b_I_*/*b_N_* < 0 and *b_I_*/*b_N_* > 0, yield qualitatively similar patterns of relative cline magnitudes between native and introduced ranges, except for defense traits; SI Fig. S2). The model included random effects of experiment identity, nested within the study identity, accounting for within- and between-study heterogeneity. Species identity was not included in the model because the multivariate analysis revealed little heterogeneity explained by species identity (analyses with species identity included did not change the results). Following the same method as before, we examined how trait category, generations-since-introduction and their interaction, gradient type, gradient overlap, hemisphere contrast, and mating system, affected the extent of parallelism between native and introduced ranges (SI Table S9). The package ‘orchaRd’ was used for graphical purposes [59].

We also conducted the same univariate analysis of estimated slopes using a composite trait value extracted using a principal component analysis (PCA) within each experiment. We conducted a PCA using *princomp* function from the package ‘stats’, with a dataset including a full set of all traits investigated by authors. The cline slope for the first principal component (trait PC1) against environmental gradients was estimated for native and introduced ranges as above.

## Results

### Multivariate analysis of clinal divergence

We found that the alignment of native and introduced range vectors (cos(*θ*)) was positive in 46 of 78 (59.0%) of experiments. The mean alignment between native and introduced cline vectors was 0.30 ([0.13, 0.46], *t_73_* = 3.6, *P* = 0.0005), which indicates partial alignment between native and introduced clines. The average log ratio of cline vector magnitudes (the mean of ln(‖***b_I_***‖/‖***b_N_***‖)) was -0.27 ([-0.52, -0.016], *t_73_* = 2.1, *P* = 0.037, Fig. 2A), which indicates that clines in introduced ranges tend to be shallower than native range clines (76.5% of native cline magnitudes, on average). Residual heterogeneity (i.e., QE) was significant, primarily due to study identity in both metrics (SI Table S3).

**Figure 2.**
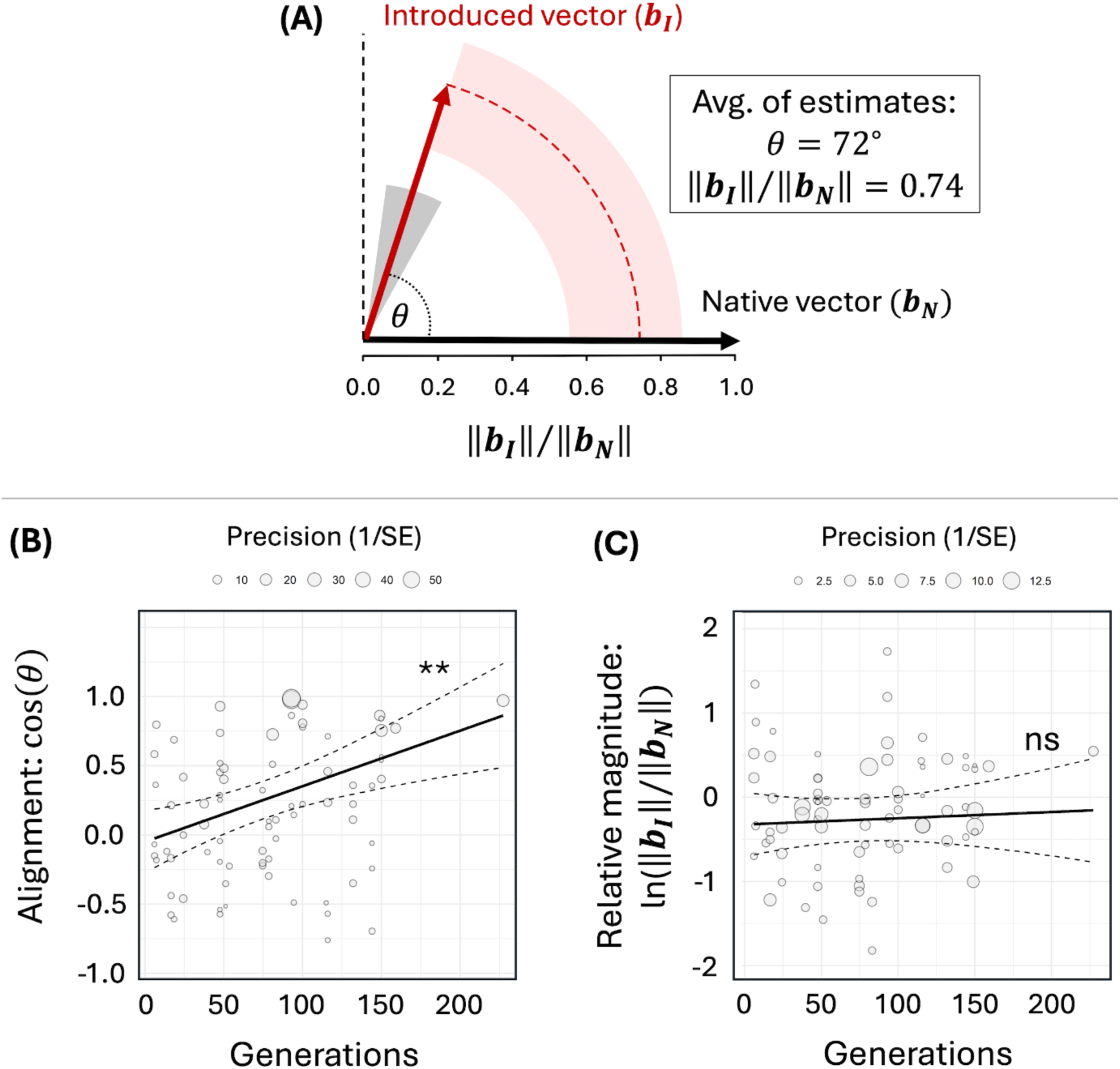
Multivariate clinal divergence between introduced and native ranges. **Panel A** shows cline vectors of native and introduced ranges presented as arrows (black and red, respectively), with the angle between the vectors (*θ*) indicating the mean degree of divergence between the orientations of native and introduced trait cline vectors. The length of the introduced vector represents the mean magnitude of introduced vector relative to that of the native vector (‖***b_I_***‖/‖***b_N_***‖), where the native vector magnitude is scaled to one (‖***b_N_***‖ = 1). Grey shading denotes the 95% CI of *θ*, and pink shading denotes the 95% CI of ‖***b_I_***‖/‖***b_N_***‖. **Panels B and C** show associations between the number of generations since introduction and **(B)** cos(*θ*) or **(C)** ln(‖***b_I_***‖/‖***b_N_***‖), with meta-analytic data shown as bubble plots. Asterisks indicate a significant association with time (** *P* < 0.001).

Variation in vector alignment was best explained by the effect of generations since introduction and the signal-to-noise ratio (*S:N*) (SI Table S4A), with cos(*θ*) increasing with introduction time (*slope* = 0.0040, *t_74_* = 3.5, *P* = 0.0008; this corresponds to a rate of increase of 0.004 per generation for cos(*θ*); Fig. 2B) and with ln(*S*: *N*) (*slope* = 0.18, *t_74_* = 4.2, *P* < 0.0001, SI Fig. S3). None of the other moderator variables were associated with cos(*θ*) (SI Table S4C). In contrast, ln(‖***b_I_***‖/‖***b_N_***‖) did not correlate with generations since introduction (*slope* = 0.0007, *t_74_* = 0.4, *P* = 0.69, Fig. 2C) or any of the other moderators (SI Table S5A). These results are robust to the removal of a single study with an exceptionally large number of generations since introduction (> 200 generations); the positive association between cos(*θ*) and time since introduction remained after the putative outlier was removed (*slope* = 0.0034, *t_73_* = 2.7, *P* = 0.009, SI Fig. S4, SI Table S4B&D), and there remains no association between ln(‖***b_I_***‖/‖***b_N_***‖) and time (SI Table S5B). These results indicate an increase in parallelism with time since introduction that is primarily due to improved alignment of cline orientations rather than changes in the relative magnitudes of multivariate clinal divergence between ranges. Since our analysis controls for variation in *S:N*, the positive association we observe between cos(*θ*) and the time since introduction appears to be a genuine signal of increased parallelism in old relative to recent introductions.

The results summarized above are for clines associated with the spatial or environmental gradients chosen by authors of the original studies. To evaluate the robustness of these results, we repeated the multivariate analyses using re-estimated trait clines predicted by absolute latitude, or gradients in temperature or precipitation (full statistical output for these follow-up tests, including p-values, are provided in SI Tables S6-S7). Results for cos(*θ*) and ‖***b_I_***‖⁄‖***b_N_***‖ were similar for each predictor of clinal divergence, though clines across precipitation gradients resulted in quantitatively weaker degrees of parallelism relative to other gradients (SI Table S6). Associations with time since introduction remained significantly positive for latitudinal clines, but not for clines predicted by temperature or precipitation gradients (SI Table S7).

### Univariate analysis of divergence across different trait categories

Among single trait cline estimates, roughly half (270 out of 513 traits, or ∼53%) showed a statistically significant cline in at least one range (Fig. 3). Among clines that were significant in one range but not the other, significant native-range clines were 2.6 times more common than significant introduced-range clines (151 vs. 58), which implies that clines are more often lost than gained in introduced ranges.

**Figure 3.**
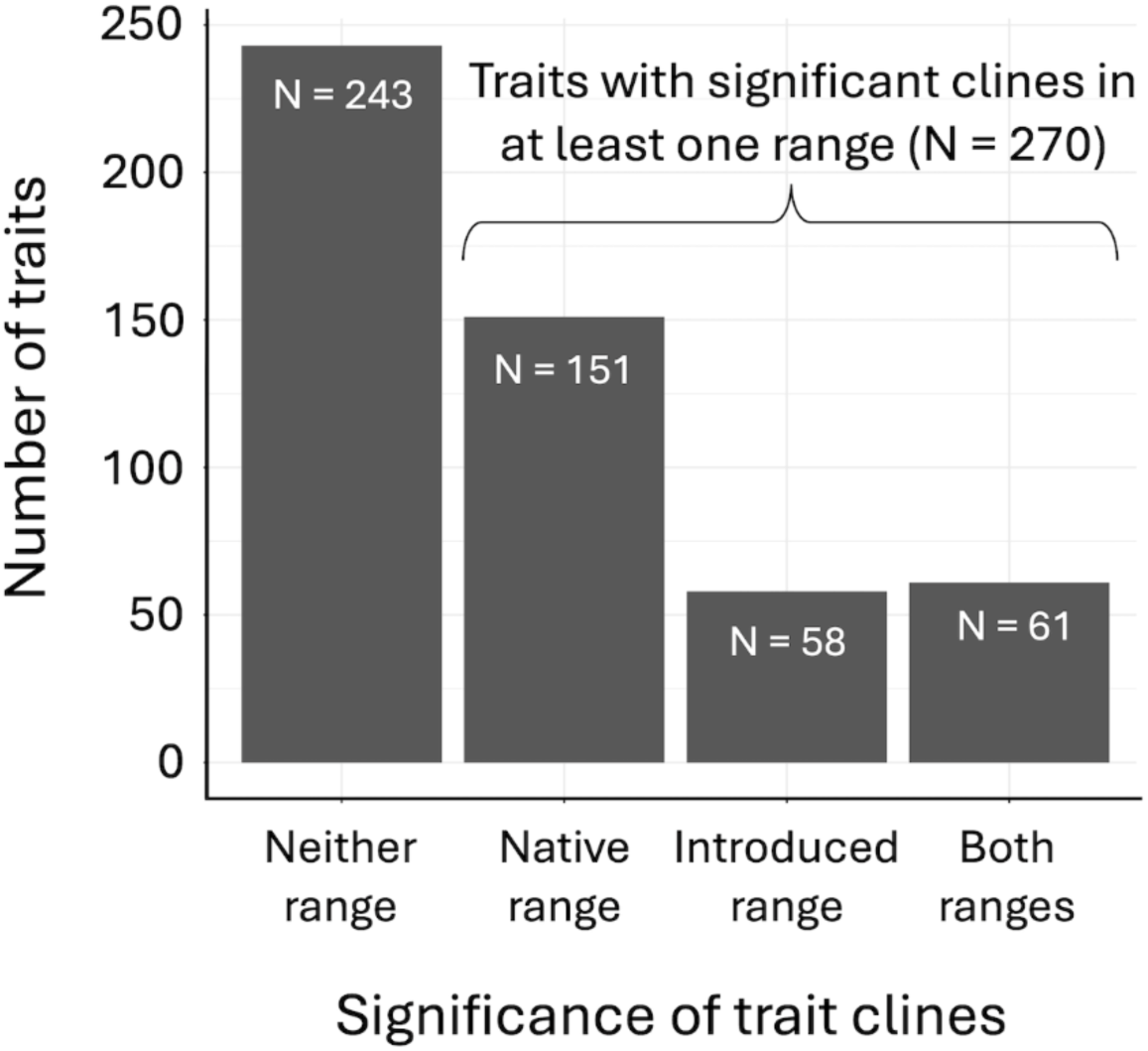
Results of the linear model tests for clinal variation in 513 traits of the meta-analysis. Slope estimates were (left to right): non-significant in both ranges, significant in the native range only, significant in the introduced range only, or significant in both ranges.

Among the 270 trait pairs with a significant cline in at least one range (Fig. 3), 59% (160 of 270) of the point estimates diverged in the same direction in each range (*b_I_*/*b_N_* > 0) and 41% diverged in opposite directions (*b_I_*/*b_N_* < 0). The proportion of cline estimates that diverged in the same direction increased with the number of generations since introduction (*slope* = 0.010, z = 3.7, *P* = 0.0002; see Fig. 4A). This pattern holds qualitatively for each trait category and trait PC1 analyzed separately, though interactions with time were only significant for the size-related traits, and marginally significant for phenology traits (SI Fig. S5), possibly owing to the smaller sample sizes of other trait categories.

**Figure 4.**
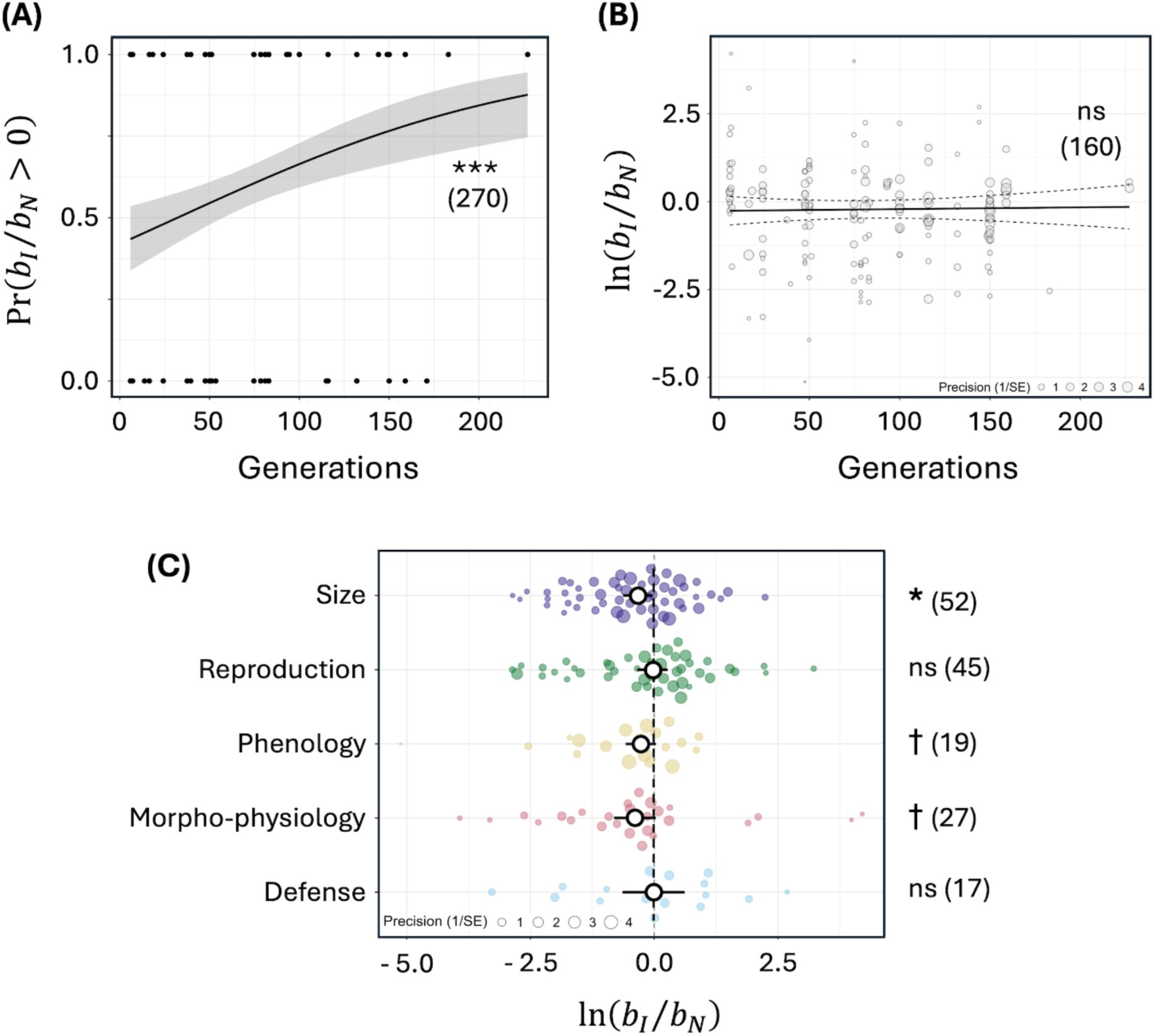
Univariate estimates of parallelism between native and introduced ranges. **Panels A-B** show the relationship between the number of generations since introduction and **(A)** the probability that clines in each range diverge in the same direction (i.e., *b_I_*/*b_N_* > 0, analyzed using a generalized linear model), or **(B)** the relative magnitude of cline slopes in each range (ln(*b_I_*/*b_N_*), calculated for traits where *b_I_*/*b_N_* > 0). **Panel C** shows ln(*b_I_*/*b_N_*) values for the same set of traits as in **(B)** but analyzed separately for each trait category. Mean effect sizes in **(C)** (white circles) are plotted with 95% confidence intervals (bold line), with the number of traits given in brackets (SI Table S7). Asterisks indicate significant deviations from ln(*b_I_*/*b_N_*) = 0 (perfect parallelism). Individual effect sizes (colored circles) for the five trait categories are plotted with circle size denoting precision (inverse variance). Asterisks indicate a significant association with time. † *P* < 0.1, * *P* < 0.05.

For the subset of cline slope estimates with the same sign (those with *b_I_*/*b_N_* > 0), native clines were similar to introduced clines, with the average magnitude of introduced clines being 80% of their native counterparts (mean ln(*b_I_*/*b_N_*) = -0.22, 95% CI = [-0.47, 0.02]). Residual heterogeneity (i.e., QE) was significant, primarily due to study identity (SI Table S3). Variability in ln(*b_I_*/*b_N_*) was not associated with the time since introduction (*slope* = -0.0005, t = 0.2385, *P* = 0.81; Fig. 4B). The magnitude of clinal divergence was similar between ranges for defense and reproductive traits, with introduced clines showing 99% and 97% the magnitude of native clines, respectively (mean ln(*b_I_*/*b_N_*) = -0.01 [-0.64, 0.61] and ln(*b_I_*/*b_N_*) = -0.04 [-0.34, 0.27], SI Table S8). In contrast, introduced clines were marginally shallower than native clines for morpho-physiology (68%, ln(*b_I_*/*b_N_*) = -0.39 [-0.82, 0.03]), and phenology (76%, ln(*b_I_*/*b_N_*) = -0.27 [-0.58, 0.04]). A subset of phenology traits (n = 15) that are specifically associated with flowering timing also showed marginally shallower introduced clines than native clines (74%, ln(*b_I_*/*b_N_*) = -0.30 [-0.61, 0.01]). Significantly shallower clines in introduced ranges were found for size traits only (72%, ln(*b_I_*/*b_N_*) = -0.33 [-0.63, -0.03]) (Fig. 4C). The magnitude of divergence in trait PC1 was similar between ranges, with introduced clines showing 94% of native clines (mean ln(*b_I_*/*b_N_*) = -0.06 [-0.42, 0.29]). We used model selection based on AICc (*glmulti*) to reduce the full model (SI Table S9) and identified the null model as the best-supported model. Although three single-predictor models were within 2 AICc units of the top model, all predictors had low summed Akaike weights (<0.4), indicating weak support for any predictor explaining variation in ln(*b_I_*/*b_N_*).

## Discussion

Our meta-analysis reveals three broad patterns of evolutionary parallelism between the trait clines of native versus introduced ranges. First, multivariate parallelism across the entire dataset is relatively low, on average, owing to shallower magnitudes of clinal divergence in introduced relative to native ranges (mean ‖***b_I_***‖⁄‖***b_N_***‖ ≈ 0.76) and modest alignment of cline directions between ranges (mean cos(*θ*) ≈ 0.30). Second, multivariate parallelism between ranges increases with the estimated residence time of introduced populations (generations since introduction), resulting in stronger parallelism between ranges involving relatively old introductions (i.e., the linear model predicts that both ‖***b_I_***‖⁄‖***b_N_***‖ and cos(*θ*) increase with time since the introduction, though the increase in ‖***b_I_***‖⁄‖***b_N_***‖ is not statistically significant; see Fig. 2B). This increase in parallelism over time is primarily caused by increased alignment of cline directions between ranges (i.e., an increase in cos(*θ*) and a declining proportion of counter-clines in the univariate data) rather than changes in the relative magnitudes of clines. Third, degrees of parallelism between ranges vary among traits, with cline slope estimates showing similar magnitudes between ranges for some categories (reproductive and defense traits) and shallower introduced-range magnitudes for others (phenological, morpho-physiological, and size-related traits, though only the latter exhibit strong statistical support for weaker clines in introduced ranges). We note that there is extensive variability among studies with respect to parallelism and the tempo with which it arises, with some studies showing strong parallelism between ranges within relatively few generations since introduction, and others showing weak parallelism more than a century after the introduction. Below, we discuss processes that might explain these observations before outlining opportunities for future work.

### Factors affecting the relative magnitudes of trait clines in native and introduced ranges

Several evolutionary mechanisms could contribute to shallower clines in introduced relative to native ranges (*i.e.*, ‖***b_I_***‖⁄‖***b_N_***‖ < 1). Most notably, shallower clines in introduced ranges are expected when there are time lags in local adaptation and introduced populations are still in the process of adapting to environmental conditions across their new ranges [60]. On the one hand, the time-lag hypothesis predicts that the ratio of cline magnitude in introduced versus native ranges (‖***b_I_***‖⁄‖***b_N_***‖) should increase with the residence time of introduced populations, yet our compiled estimates of ‖***b_I_***‖⁄‖***b_N_***‖ showed, at best, a weak (and non-significant) increase with residence time (see Fig. 2C). Similarly weak interactions with time were observed for univariate clines of different trait categories. On the other hand, such patterns are expected if trait clines arise rapidly during the range expansion itself, such that later changes in cline magnitudes are moderate relative to the rates at which they initially arise.

Clines can emerge during the initial establishment and geographic spread of invasive species by several mechanisms, including: (1) adaptation during the range expansion, (2) preferential establishment of locally pre-adapted populations across the new range [33,61], (3) population structure caused by multiple independent introductions across the new range, followed by incomplete mixing when these populations come into contact [34], and (4) strong genetic drift owing to serial bottlenecks during geographic expansion within the new range [22]. Initially rapid rates of cline formation may be difficult to detect in datasets like ours unless there is fine-scale temporal resolution across many species with very recent introductions. Further work is needed to test the extent to which adaptive versus nonadaptive clines arise in recent introductions, through experiments that control for effects of genetic ancestry from multiple introductions [34], and tests for alignment between clinal divergence and the directions of local selection at different points across the range. Time-series data (e.g., from herbarium specimens) may also show that early mismatches between ranges are later resolved as introduced populations evolve clines that better match native clines (e.g., the hb-chr5b haplotype in common ragweed [28]). Similarly, spatial sorting of dispersal-related traits during expansion, documented in both plants and animals (see [62,63]), can create gradients that are potentially correlated with environmental variables but primarily reflect the history of range expansion rather than local adaptation.

Differences between ranges in the steepness of environmental gradients represent an additional factor potentially causing differences in native and introduced trait clines. Shallower clines in the introduced range may be expected, even at equilibrium, in cases where environmental gradients are shallower in the introduced compared to the native range. Trait clines might be shallower, on average, in introduced ranges if species are disproportionately introduced into geographic regions where the environmental gradients are relatively shallow, which would facilitate range expansions in those regions [30]. We evaluated this hypothesis by comparing the steepness of environmental gradients for temperature and precipitation between native and introduced ranges (i.e., the rate at which these variables change per unit of distance in latitude or elevation). We found no systematic differences between the ranges, suggesting that the overall shallower clines in the introduced ranges are due to other factors (see SI Table S10 for details).

A more complex picture emerges in the presence of genetic, evolutionary, and demographic factors that constrain local adaptation, potentially leading to shallower clines in introduced ranges (see SI Appendix 2). For example, traits under weaker selection or with lower genetic variation in the introduced range may evolve shallower clines at migration-selection balance, owing to stronger effects of swamping by gene flow. Previous work suggests that heritability estimates [64–68] and linear selection gradients [22] do not systematically differ between native and introduced ranges, which offers little support for this possibility, though such data are admittedly sparse and the hypothesis deserves further attention. Demographic differences between ranges—including differences in the rate of dispersal or the spatial distribution of population density (see SI Appendix 2)—can also lead to different degrees of swamping of local adaptation between ranges. Invasive herbaceous plant species tend to have higher dispersal rates than non-invasive ones [69], and increased dispersal ability can evolve within introduced ranges [64,70], which could lead to higher swamping by gene flow in introduced ranges relative to native ranges. The distribution of population density affects local adaptation by mediating the rate of gene flow from the center of a range to its edges (see SI Appendix 2; [29]). If, for example, introduced range populations more closely conform to an “abundant-center” distribution than native range populations (which might be likely in actively expanding introductions; [71]) they will be more susceptible to swamping and evolve shallower clines.

The trend towards shallower clines in introduced ranges does not, however, apply uniformly. Defense and reproductive traits had similar cline magnitudes between ranges, while cline estimates for phenological, morpho-physiological, and size-related traits were shallower in introduced relative to native ranges (introduced clines are, on average, 68-76% as steep as native clines, with only size-related traits showing a significant difference between ranges). Variation among traits might stem from differences in evolutionary potential for rapid local adaptation, with traits under exceptionally strong selection or with high heritabilities evolving more rapidly in introduced ranges. Reproductive traits, for example, are thought to be under strong selection because they are closely related to fitness [72,73]. Likewise, the average heritability of plant secondary metabolites (which includes defensive compounds) is estimated to be more than twice as large as the heritability of phenological, morphological, physiological, and size traits, which should facilitate strong responses to selection ([74]; see also [64,75]).

### Factors affecting the alignment of cline directions between ranges over time

Three factors potentially contribute to the weaker alignment of cline directions in more recent than older introductions (recall that cos(*θ*) quantifies alignment between ranges). First, traits differ in how responsive they are to natural selection: those with high heritability and under strong selection will rapidly adapt to local conditions across the new range, while those with low heritability or weak selection should exhibit protracted lags in local adaptation (see the SI Appendix 2). This heterogeneity of lag-times among traits leads to temporary misalignments in the relative magnitudes of clines in each range, which eventually resolve as each trait evolves to migration-selection equilibrium in the new range (see the black broken curve in Fig. 1C).

Second, maladaptive clines that emerge at the outset of a range expansion can strongly depress the initial alignment of clinal divergence between ranges. For example, if introduced clines initially arise by genetic drift during the range expansion, then the alignment of clines between ranges will initially be poor (cos(*θ*) ≈ 0 initially, on average) with a large proportion of traits showing counter-clines (Fig. 1C). If selection pressures are similar between ranges, stochastic and historical effects on the initial formation of clines are expected to diminish over time, as natural selection reorients clines to align with those that are selectively maintained in the native range, leading to the observed increase in parallelism among the older introductions. Strong genetic correlations between traits can also generate transient counter-clines in traits subject to relatively weak selection (see SI Appendix 6). Strong parallelism of clines (e.g., ‖***b_I_***‖⁄‖***b_N_***‖ ≈ cos(*θ*) ≈ 1) can ultimately evolve between the native and introduced ranges, yet strong genetic constraints can draw out this process, leading to extended lags in local adaptation of introduced populations and extensive changes over time in the alignment of clines between ranges. Strong genetic correlations and trade-offs between fitness-related traits may indeed be common. For example, both early flowering and large biomass (and seed production) are favored at high latitudes, yet these traits negatively covary, providing the potential for the evolution of transient counter-clines in the more weakly selected trait (see [20,76]).

Third, the directions of local selection across each range might change over time, leading to a greater alignment of both selection and trait clines in old relative to recent introductions. For example, selection may fundamentally differ between recent and old introductions if the former experiences a release from natural enemies, but the latter no longer does. Enemy release might relax selection of defense traits within the new range (as predicted by the “evolution of increased competitive ability” (EICA) hypothesis [77]), and this could facilitate evolution of defense trait clines caused by drift or by indirect responses to selection of genetically correlated traits. Moreover, relaxation of herbivory may alter the pattern of selection on secondary metabolites with dual functions in herbivore defense and abiotic stress tolerance [21], with metabolite concentrations in recently introduced species evolving in response to selection for abiotic stress tolerance rather than herbivore defense [21]. Clines favored by selection in recent introductions may, thus, differ substantially from those in the native range. However, subsequent accumulation of natural enemies in the introduced range over time [78,79] may gradually reshape the selective landscapes of older introduced ranges, leading to a stronger alignment of selection and evolutionary parallelism of clines between native and introduced ranges.

### Conclusions and future directions

A trait or allele frequency cline is not, in itself, evidence for local adaptation [22]. However, repeated evolution of parallel clines between geographically disjunct regions of a species’ range is difficult to reconcile with nonadaptive explanations and is instead suggestive of repeated local adaptation to shared environmental conditions between the regions [16,17]. We found that geographic clines in plant traits show strong evolutionary parallelism between native and introduced ranges with relatively long residence times (on the order of 100 or more generations). This result applies to a broad set of plant species and trait types and implies that local adaptation is extensive and predictable across native ranges and mature introduced ranges.

Our analysis suggests that the greater parallelism of older relative to more recent introductions is primarily due to an increase in the alignment of cline directions between ranges rather than changes to the relative magnitudes of clines in introduced versus native ranges—a pattern that holds qualitatively across different trait categories. Although several factors may contribute to this pattern, we suspect it is caused by rapid emergence of maladaptive clines in recent introductions—owing to processes like drift, founder effects, and spatial sorting during colonization (see Fig. 1C), or to indirect responses to selection of genetically correlated traits (see SI Appendix 6). This early stage of cline formation may then be followed, in the longer term, by more gradual evolution of local adaptation that increases parallelism between clines in each range. Direct tests for enrichment of maladaptive clines in recent relative to old introductions (e.g., through estimates of selection of clinally diverging traits) would, therefore, be a worthwhile focus for future work. Other potentially useful extensions of our study include analyses of nonlinear clines (which are sometimes reported in cline studies, e.g., [11,80]) and their evolutionary parallelism between ranges, and analyses of the joint tempo of changes in trait clines and the geographic breadths of introduced species’ ranges, as adaptive clines can be shaped by and facilitate species’ range breadth [30].

Despite these patterns, we emphasize that there is extensive variation in metrics of cline parallelism that are not explained by covariates of our meta-analysis models, which included plant mating systems, whether ranges are confined to the same hemisphere, and the similarity of abiotic environmental gradients between ranges. Much of the unexplained variability could be attributable to other features of the species or ranges that they span, including similarity of biotic conditions between native and introduced ranges. Future work is required to better understand why patterns of parallelism vary so widely, and whether similar patterns apply to allele frequency clines.

Finally, we used a snapshot of clinal divergence in species that differed in their arrival times into new ranges to indirectly infer temporal dynamics of cline evolution. However, a true time-series analysis of evolutionary parallelism between native and introduced clines would also be valuable, though such data are much more difficult to obtain. In a rare exception, Wu and Colautti [20] analyzed herbarium records to compare clinal patterns of flowering phenology in three introduced ranges of *Lythrum salicaria* over 150 years, and found lags in the evolution of adaptive clines in all three regions. Because herbarium specimens are collected in the field, the contribution of phenotypic plasticity to the trait variation must be controlled [20], although genomic analyses [28,81] or quantitative genetic approaches can help overcome those issues [82]. It will be interesting to see how strongly true time-series analyses conform to the patterns of increasing parallelism implied by our study.

## Acknowledgements

We are grateful to three anonymous reviewers, whose thoughtful comments on an earlier version of the paper led to substantial improvements in the analysis, presentation, and interpretation of the results. This research was supported by funds from the Australian Research Council (DP220102706 to AU, and DP240102637 to KH, TC, and AU), RMIT, and Monash University.

