## Supporting Information for "Meta-analysis reveals the tempo of evolutionary parallelism of local adaptation between native and introduced ranges of invasive plant species"

#### List of contents:


- **Appendix 1:** Models for the evolution of clines in quantitative traits
- **Appendix 2:** Factors leading to different trait clines between ranges
- **Appendix 3:** Multivariate clinal divergence when clines are initially zero
- **Appendix 4:** Serial bottlenecks and drift-induced clines
- **Appendix 5:** Clinal divergence when the initial clines are drift-induced
- **Appendix 6:** Effects of genetic correlations between traits
- **Supplementary References**
- **Supplementary Figures S1-S2,**
- **Box S1**
- **Supplementary Tables S1-S10**
- **Supplementary Figures S3-S5**

#### Appendix 1: Models for the evolution of clines in quantitative traits

Following García-Ramos and Kirkpatrick (1997), a cline for a quantitative trait under Gaussian selection to a local optimum can be modelled as follows. We consider a species spread over a continuous geographic range in one dimension (*e.g.*, by latitude), with $x$ representing the location within the range. Generations are discrete and non-overlapping. We assume that clines evolve deterministically, in response to gene flow and selection, following the range expansion, and once the spatial distribution of population sizes in a range becomes stable, at time $t = 0$. From this point onward, the change in the trait mean at position $x$ in the range will be:

$$\Delta\bar{z}(x,t) = \frac{\sigma^2}{2}\frac{\partial^2\bar{z}(x,t)}{\partial x^2} + \sigma^2\frac{\partial \log\big(N(x)\big)}{\partial x}\frac{\partial\bar{z}(x,t)}{\partial x} + h^2\beta(x,t) \qquad \text{(S1)}$$

where $\sigma^2$ is the variance of dispersal distances among individuals, $h^2$ is the narrow-sense heritability of the trait ($0 \leq h^2 \leq 1$), $N$ is the population density distribution across the gradient, $\beta(x,t) = \gamma\big(O(x) - \bar{z}(x,t)\big)$ is the linear selection gradient at position $x$ in the gradient, $O(x)$ is the optimal trait value at position $x$, and $\gamma$ is the strength of stabilizing selection around the optimum. Our expressions assume that the trait (at each location) is normally distributed with unit variance, so that the heritability is equivalent to the additive genetic variance (thus, the phenotypic variance is scaled to one). Under these assumptions, $\gamma = (\omega + 1)^{-1}$, where $\omega$ is the width of the Gaussian fitness function (see García-Ramos and Kirkpatrick 1997). Because $\omega$ is constrained to be positive and finite, the strength of stabilizing selection, $\gamma$, must meet the constraint: $0 < \gamma < 1$. The strength of stabilizing selection, genetic variance, residual variance, and heritability are all assumed to be constant across the gradient. In what follows, we assume, unless otherwise stated, that the optimum changes linearly across the range (*i.e.*, $O(x) = Bx$, where $B$ is the slope of the optimum change), in which case, eq. (S1) can be rewritten as follows:

$$\Delta\bar{z}(x,t) = \frac{\sigma^2}{2}\frac{d^2\bar{z}(x,t)}{dx^2} + \sigma^2\frac{d\bar{z}(x,t)}{dx}\frac{d\ln(n)}{dx} + h^2\gamma\big(Bx - \bar{z}(x,t)\big)$$

We will consider two extreme models for the distribution of population density across a range. These extremes can be thought of as bookend cases of the model, with intermediate cases between them yielding intermediate results. At one extreme, which we refer to as the ***"uniform density" model***, population density is uniformly distributed across the range. Previous work has shown that uniform density across the range will lead to relatively symmetric patterns of gene flow between centre and range edge populations, which tends to facilitate strong local adaptation to the optimum (García-Ramos and Kirkpatrick 1997).

At the other extreme, population density is normally distributed with highest density at the range centre (with no loss of generality, we let $x = 0$ represent the range centre in our analysis). This scenario, which is referred to as the ***"abundant centre" model***, leads to strong asymmetries in gene flow between centre and edge populations across the range. Dispersal from the centre to the range margins is much higher than the reverse, leading to high rates of maladaptive gene flow from the centre to the edge, which inhibits local adaptation. Consequently, trait clines tend to be much shallower at equilibrium in abundant centre models relative to uniform density models (García-Ramos and Kirkpatrick 1997).

**Evolution of trait clines under a model of uniform population density**

Under the uniform density model, with abrupt range boundaries at positions $x_{\text{min}} = -x^*$ and $x_{\text{max}} = x^*$, eq. (S1) simplifies to:

$$\Delta\bar{z}(x,t) = \frac{\sigma^2}{2}\frac{\partial^2\bar{z}(x,t)}{\partial x^2} + h^2\gamma\big(Bx - \bar{z}(x,t)\big) \tag{S2}$$

Assuming migrants do not disperse beyond the range boundaries (*i.e.*, the range boundaries are "reflecting") eq. (S2) has the following equilibrium solution:

$$\bar{z}_{eq.}(x) = Bx + B\frac{\exp\left((x^* - x)\sqrt{\frac{2h^2\gamma}{\sigma^2}}\right) - \exp\left((x^* + x)\sqrt{\frac{2h^2\gamma}{\sigma^2}}\right)}{\sqrt{\frac{2h^2\gamma}{\sigma^2}}\left(1 + \exp\left(2x^*\sqrt{\frac{2h^2\gamma}{\sigma^2}}\right)\right)} \tag{S3}$$

(see eq. (12) of García-Ramos and Kirkpatrick 1997). When selection and heritability are strong relative to gene flow (in which case $2h^2\gamma \gg \sigma^2$), or the range is sufficiently broad that $x^*\sqrt{\frac{2h^2\gamma}{\sigma^2}} \gg 1$, then the second term of the equilibrium in eq. (S3) tends to be small, and the slope of the trait cline at equilibrium is approximately $b_{eq.} = d\bar{z}_{eq.}(x)/dx \approx B$ across most of the range, though this approximation is inaccurate near the range edges, where the slope of the trait cline becomes somewhat shallower than predicted by the approximation; see Fig. 1A, which presents plots of the exact equilibrium based on eq. (S3)).

The intermediate dynamics of the cline can be approximated by assuming an initial condition in which the trait cline is either linear or absent ($\frac{\partial^2 \bar{z}(x,t)}{\partial x^2} = 0$, in either case), and that edge effects due to the "reflecting boundaries" assumption are small and largely localized to the range margins. Under such conditions, then the difference equation simplifies to:

$$\Delta \bar{z}(x,t) = h^2 \gamma \big(Bx - \bar{z}(x,t)\big) \tag{S4}$$

The general solution is then:

$$\bar{z}(x,t) = Bx - \big(Bx - \bar{z}(x,0)\big)(1 - h^2\gamma)^t \tag{S5}$$

where $\bar{z}(x,0)$ is the initial trait mean at location *x* in the introduced range and $\bar{z}(x,t)$ is the trait mean after *t* generations. The cline slope at time *t* is thus:

$$b(t) = \frac{\bar{z}(x,t)}{dx} = B(1 - (1 - h^2\gamma)^t) + b_0(1 - h^2\gamma)^t \tag{S6}$$

where $b_0 = \frac{d\bar{z}(x,0)}{dx}$ is the cline slope at time *t* = 0. Note that the result depends on the initial value of the cline slope ($b_0$), but not its intercept, defined as the initial mean expressed at the range centre (i.e., $\bar{z}(0,0)$). Note that edge effects (the "reflecting boundaries" in García-Ramos and Kirkpatrick 1997) are expected to result in departures from cline linearity near the range margins, though such effects are trivial over most of the range when selection is strong relative to dispersal.

**Evolution of trait clines under an abundant center model of population density**

Now consider the case where the population density distribution conforms to an "abundant center" model. Let the population density as a function of location in the range follow a Gaussian function with density maximized at the center of the range:

$$N(x) = N_{\max} \exp\left(-\frac{1}{2}x^2\right) \tag{S7}$$

in which case $\frac{\partial \log(N(x))}{\partial x} = -x$, and eq. (S1) simplifies to:

$$\Delta \bar{z}(x,t) = \frac{\sigma^2}{2}\frac{\partial^2 \bar{z}(x,t)}{\partial x^2} - \sigma^2 x \frac{\partial \bar{z}(x,t)}{\partial x} + h^2\gamma\big(Bx - \bar{z}(x,t)\big) \tag{S8}$$

If the cline is linear or absent at an arbitrary time *t* (thus, $\bar{z}(x,t) = xb_t$, $\frac{\partial^2 \bar{z}(x,t)}{\partial x^2} = 0$ and $\frac{\partial \bar{z}(x,t)}{\partial x} = b_t$), then the trait mean at the next generation will remain linear. This can be seen from the recursion equation for the trait mean at position *x* in the range, which becomes:

$$\begin{aligned}\bar{z}(x,t+1) &= \bar{z}(x,t) - \sigma^2 x \frac{\partial \bar{z}(x,t)}{\partial x} + h^2\gamma\big(Bx - \bar{z}(x,t)\big) \\ &= x\big(b_t - \sigma^2 b_t + h^2\gamma(B - b_t)\big) \qquad \text{(S9)}\end{aligned}$$

Thus, if the cline is initially linear, then it will remain linear under the abundant center model for population density.

If the cline is initially linear or absent, we can substitute $\bar{z}(x,t) = c(t) + \frac{d\bar{z}(x,t)}{dx}x$, where $c(t) = \bar{z}(0,t)$ is the intercept at time *t*. The intercept in the next general is then:

$$c(t+1) = c(t)(1 - h^2\gamma) \qquad \text{(S10)}$$

The recursion can be rewritten as:

$$\bar{z}(x,t+1) = c(t+1) + x\left(\frac{d\bar{z}(x,t)}{dx} - \sigma^2\frac{d\bar{z}(x,t)}{dx} + h^2\gamma\left(B - \frac{d\bar{z}(x,t)}{dx}\right)\right) \qquad \text{(S11)}$$

Substituting $\bar{z}(x,t+1) = c(t+1) + \frac{d\bar{z}(x,t+1)}{dx}x$, we have:

$$\frac{d\bar{z}(x,t+1)}{dx} = \frac{d\bar{z}(x,t)}{dx} - \sigma^2\frac{d\bar{z}(x,t)}{dx} + h^2\gamma\left(B - \frac{d\bar{z}(x,t)}{dx}\right) \qquad \text{(S12)}$$

which is notably independent of the intercept among the introduced populations, as $c(t+1)$ and $c(t)$ drop out of the recursion. The recursion for the slope of the cline ($b(t) = \frac{d\bar{z}(x,t)}{dx}$) is:

$$b(t+1) = b(t) - \sigma^2 b(t) + h^2\gamma\big(B - b(t)\big) \qquad \text{(S13)}$$

which has the equilibrium:

$$b_{eq.} = \frac{h^2\gamma B}{\sigma^2 + h^2\gamma} \qquad \text{(S14)}$$

(this is the equivalent of eq. (6) in García-Ramos and Kirkpatrick 1997). The general solution for the cline slope is:

$$b(t) = b_{eq.}(1 - (1 - \sigma^2 - h^2\gamma)^t) + b_0(1 - \sigma^2 - h^2\gamma)^t \qquad \text{(S15)}$$

where $b_0$ is the initial slope. Once again, the result depends on the initial value of the cline slope ($b_0$), but not its intercept. In cases where the initial slope is zero ($b_0 = 0$), we obtain the same result as we had before.

#### Appendix 2: Factors leading to different trait clines between ranges

Since the scales of measurement can differ widely among traits and species (e.g., scales over which dispersal occurs), it is useful to present comparisons of clines in introduced and native range populations by estimating the ratios of the trait cline slopes in one range relative to the other. In our discussion of cline theory and the meta-analysis, we focus on the ratio $b_I/b_N$, which is expected to be 1 whenever the clines of a trait shift at the same rate and direction in both ranges (i.e., $b_I = b_N$, regardless of the absolute magnitudes of $b_I$ and $b_N$). In this section, we use the cline theory models outlined above to describe the set of conditions that can cause introduced and native trait clines to differ (i.e., conditions leading to $b_I \neq b_N$).

Under the stated assumptions of the trait cline models presented above, there are six specific scenarios that can cause trait clines to differ between native and introduced ranges of a species. These differences, which we deal with in turn, include:

1. ***Evolutionary lags in the introduced range***, owing to the relatively limited time for local adaptation to have evolved in the introduced range relative to the native range.
2. ***Differences between ranges in the steepness of environmental gradients***, leading to different rates of change of trait optima across the introduced range relative to the native range (*i.e.*, $B_I \neq B_N$ where $B_I$ and $B_N$ represent the rates of change of the trait optima across the introduced and native ranges, respectively)
3. ***Differences between ranges in trait heritability*** (*i.e.*, $h_I^2 \neq h_N^2$ where $h_I^2$ and $h_N^2$ are the heritabilities of the same trait in introduced and native range populations).
4. ***Different strengths of stabilizing selection between ranges*** (*i.e.*, $\gamma_I \neq \gamma_N$ where $\gamma_I$ and $\gamma_N$ represent the strengths of stabilizing selection for a trait in the introduced and native range populations, respectively).
5. ***Different rates of dispersal (gene flow) between ranges*** (*i.e.*, $\sigma_I^2 \neq \sigma_N^2$ where $\sigma_I^2$ and $\sigma_N^2$ represent the range-specific variances of dispersal distances between individuals from their birthplaces).
6. ***Different spatial distributions of population density between ranges*** (*i.e.*, different $N(x)$ functions between the introduced and native ranges).

##### 1. Evolutionary lags in the introduced range

If population density is uniformly distributed across both the introduced and the native ranges, the native range is at equilibrium, and there is initially no cline in the introduced range (at time $t = 0$), then the ratio of the trait cline slopes between the ranges, over time, will be:

$$\frac{b_I}{b_N} = \frac{b_{I,eq.}}{b_{N,eq.}}(1-(1-h_I^2\gamma_I)^t) = \frac{B_I}{B_N}(1-(1-h_I^2\gamma_I)^t) \tag{S16}$$

Provided there is some degree of selection for local adaptation across the native range ($B_N \neq 0$), then eq. (S16) implies that $b_I/b_N \approx 0$ as $t$ approaches zero (i.e., immediately after expansion into the introduced range), and that $b_I/b_N \approx B_I/B_N$ as $t$ becomes large as the introduced range approaches equilibrium.

If population density follows an abundant center distribution in each range, and there is initially no introduced cline at time $t = 0$, then the ratio of the introduced to the native cline, following $t$ generations of evolution in the introduced range, will be:

$$\frac{b_I}{b_N} = \frac{b_{I,eq.}}{b_{N,eq.}}(1-(1-\sigma_I^2-h_I^2\gamma_I)^t) \tag{S17}$$

where $b_{I,eq.} = \frac{h_I^2\gamma_I B_I}{\sigma_I^2+h_I^2\gamma_I}$ and $b_{N,eq.} = \frac{h_N^2\gamma_N B_N}{\sigma_N^2+h_N^2\gamma_N}$ are the respective equilibria for introduced and native clines slopes. Note that eq. (S17) is approximately equal to eq. (S16) when the product of heritability and the strength of selection is large relative to the dispersal rate ($h_I^2\gamma_I \gg \sigma_I^2$). More generally, $b_I/b_N$ is again expected to approach zero as $t$ approaches zero. And at the other extreme, the equilibrium (large-$t$) ratio of cline slopes approaches:

$$\lim_{t\to\infty}\frac{b_I}{b_N} = \frac{b_{I,eq.}}{b_{N,eq.}} = \frac{\frac{h_I^2\gamma_I B_I}{\sigma_I^2+h_I^2\gamma_I}}{\frac{h_N^2\gamma_N B_N}{\sigma_N^2+h_N^2\gamma_N}} \tag{S18}$$

In the special case where the trait heritability, strength of stabilizing selection, and dispersal rate, are all equal between the introduced and native ranges ($h_I^2 = h_N^2$, $\gamma_I = \gamma_N$, and $\sigma_I^2 = \sigma_N^2$), then eq. (S18) simplifies to $\lim_{t\to\infty} b_I/b_N = B_I/B_N$, which matches the long-term prediction under the uniform population density model.

Eqs. (S16-S17) imply that evolutionary lags to equilibrium in the introduced range will decay geometrically with a rate of $h_I^2\gamma_I$ in the uniform density model and a rate of $\sigma_I^2 + h_I^2\gamma_I$ in the abundant center model. In both cases, rates of decay should be rapid, and evolutionary lags highly transient, for traits under strong selection with high heritability (i.e., traits for which the compound parameter $h_I^2\gamma_I$ is relatively large). Dispersal (gene flow) further affects the timescale of the lag in the abundant center model because it causes equilibrium clines to become shallow over the entire range (in contrast dispersal typically has a more localized effect, near the range margins, in the uniform density model), and it takes less time to evolve a shallow cline.

**2. Differences between ranges in the steepness of environmental gradients**

When rates of change for the trait's optimum differ between ranges, but all else is equal between ranges, then the ratios in the uniform density and abundant center density models become (eq. (S16) and (S17), respectively, each evaluated with $b_0 = 0$):

$$\frac{B_I}{B_N}(1-(1-h_I^2\gamma_I)^t) \qquad \text{(S19)}$$

and

$$\frac{B_I}{B_N}(1-(1-\sigma_I^2-h_I^2\gamma_I)^t) \qquad \text{(S20)}$$

both of which converge to $B_I/B_N$ over time (sufficiently large *t*). In the long run, native-range clines will be steeper than their introduced-range counterparts when the environmental gradients mediating local selection are steeper in the native range than in the introduced range (i.e., $-1 < b_I/b_N < 1$ occurs when $|B_N| > |B_I|$), and introduced range clines will be steeper when introduced environmental gradients are steeper (i.e., $|b_I/b_N| > 1$ occurs when $|B_I| > |B_N|$).

**3. Differences between ranges in trait heritability**

In cases where the heritability of the trait is lower in the introduced range relative to the native range (i.e., due to an extreme bottleneck), ratios of trait cline slopes are affected in two ways. First, it can prolong evolutionary lags to equilibrium in the introduced range (recall that lags to equilibrium decay rapidly when $h_I^2\gamma_I$ is large but may persist for a long time when $h_I^2\gamma_I$ is small). Second, differences in heritability between ranges can affect the equilibrium ratio of cline slopes (equilibrium of $b_I/b_N$). Since this second effect tends to be weak in models with a uniform population density across the range, we focus immediately below on cases where population densities conform to an abundant center model.

Assuming that all else is equal between the ranges (i.e., $B_I = B_N$, $\gamma = \gamma_I = \gamma_N$, $\sigma^2 = \sigma_I^2 = \sigma_N^2$), and population density has an abundant center, then the equilibrium ratio of clines slopes (eq. (S18)) simplifies to:

$$\frac{b_{I,eq.}}{b_{N,eq.}} = \frac{h_I^2(h_N^2\gamma + \sigma^2)}{h_N^2(h_I^2\gamma + \sigma^2)} \tag{S21}$$

From eq. (S21), we see that when the product of heritability and selection are strong relative to dispersal ($h_N^2\gamma \gg \sigma^2$ and $h_I^2\gamma \gg \sigma^2$), then $b_{I,eq.}/b_{N,eq.} \approx 1$. In contrast, with high gene flow in an abundant center model, eq. (S21) simplifies to:

$$\frac{b_{I,eq.}}{b_{N,eq.}} \approx \frac{h_I^2}{h_N^2} \tag{S22}$$

which leads to steeper trait clines in the native range whenever $h_I^2 < h_N^2$, though the effect on $b_{I,eq.}/b_{N,eq.}$ should be small unless there is a substantial difference in the trait's heritability between the ranges.

**4. Different strengths of stabilizing selection between ranges**

Differences between ranges in the strength of stabilizing selection have the same qualitative and quantitative effects as differences in heritability (see the previous section). Relatively weak selection in the introduced range can, firstly, extend the period over which there is lag in local adaptation within the introduced range. Secondly, in cases where population density follows an abundant center model in both ranges (and assuming all else is equal between ranges, *i.e.*: $B_I = B_N$, $h^2 = h_I^2 = h_N^2$, $\sigma^2 = \sigma_I^2 = \sigma_N^2$), then the equilibrium ratio of cline slopes becomes:

$$\frac{b_{I,eq.}}{b_{N,eq.}} = \frac{\gamma_I(h^2\gamma_N + \sigma^2)}{\gamma_N(h^2\gamma_I + \sigma^2)} \tag{S23}$$

As before, $b_{I,eq.}/b_{N,eq.} \approx 1$ under weak gene flow ($h^2\gamma_N \gg \sigma^2$ and $h^2\gamma_I \gg \sigma^2$), and $b_{I,eq.}/b_{N,eq.} \approx \gamma_I/\gamma_N$ under strong gene flow ($\sigma^2 \gg h^2\gamma_N$ and $\sigma^2 \gg h^2\gamma_I$). Thus, clines in the introduced range will be much shallower than those in the native range if stabilizing selection is substantially weaker in the introduced range, and there is substantially more population density at the range center and high gene flow, leading to substantial swamping of local adaptation away from the center of the introduced range. Such a scenario should be particularly likely for traits that are important in the native range, but experienced relaxed selection in the introduced range, such as traits that function in defense of specialist herbivores that are abundant in the native range and absent from the introduced range.

**5. Different rates of dispersal (gene flow) between ranges**

Effects of differences in dispersal between ranges will primarily affect clines in species with abundant center population densities (much more modest effects apply in species with uniform distributions). Focusing on the specific case where dispersal rates differ between ranges, but the remaining parameters of local adaptation do not differ ($\gamma = \gamma_I = \gamma_N$, $B_I = B_N$, and $h^2 = h_I^2 = h_N^2$), the equilibrium ratio of cline slopes, assuming an abundant center, becomes:

$$\frac{b_{I,eq.}}{b_{N,eq.}} = \frac{\sigma_N^2 + h^2\gamma}{\sigma_I^2 + h^2\gamma} \quad \text{(S24)}$$

which simplifies to $b_{I,eq.}/b_{N,eq.} \approx 1$ when gene flow is weak ($h^2\gamma \gg \sigma_I^2, \sigma_N^2$) and $b_{I,eq.}/b_{N,eq.} \approx \sigma_N^2/\sigma_I^2$ when gene flow is strong ($\sigma_I^2, \sigma_N^2 \gg h^2\gamma$). In general, we expect steeper clines in the native range when dispersal is stronger in the introduced range ($b_{I,eq.}/b_{N,eq.} < 1$ when $\sigma_I^2 > \sigma_N^2$), and steeper clines in the introduced range when dispersal is highest in the native range ($b_{I,eq.}/b_{N,eq.} > 1$ when $\sigma_N^2 > \sigma_I^2$). Gene flow must be relatively high, leading to substantial swamping of local adaptation towards the range margins, for these effects are pronounced (e.g., $b_{I,eq.}/b_{N,eq.} \ll 1$ requires $\sigma_I^2 \gg h^2\gamma$ and abundant center population density).

**6. Different spatial distributions of population density between ranges**

Differences between ranges in the spatial distributions of population density can profoundly affect the relative opportunities for local adaptation in each range. For example, suppose the introduced range follows an abundant center distribution and the native range has a uniform distribution, and all else is equal between the ranges. The ratio of equilibrium cline slopes between the ranges will be:

$$\frac{b_{I,eq.}}{b_{N,eq.}} = \frac{B_I}{B_N} \cdot \frac{h_I^2 \gamma_I}{\sigma_I^2 + h_I^2 \gamma_I} \tag{S25}$$

Note that the term $h_I^2\gamma_I/(\sigma_I^2 + h_I^2\gamma_I)$ is constrained to be less than one and greater than zero provided the population can respond to selection ($h_I^2\gamma_I > 0$) and there is at least some gene flow across the range ($\sigma_I^2 > 0$). Consequently, trait clines will tend to be shallower in the introduced range than in the native range (for the specific scenario outlined) unless $B_I/B_N$ has a substantial enough elevation above one to offset the constraining effect of abundant center density within the introduced range. For example, in a scenario with selection and heritability of similar magnitude to dispersal ($h_I^2\gamma_I = \sigma_I^2$), the environmental gradient in the introduced range must be double the gradient in the native range for the clines to be equal (*i.e.*, $B_I/B_N = 2$ leads to $b_{I,eq.}/b_{N,eq.} = 1$). In cases where the environmental gradients are equal ($B_I/B_N = 1$), then $b_{I,eq.}/b_{N,eq.} < 1$.

#### Appendix 3: Multivariate clinal divergence when clines are initially zero

As before, we assume that the native range is at migration-selection equilibrium, while the introduced range is evolving to its equilibrium. Let us assume that there are *n* traits under spatially varying selection in each range. The native and introduced multivariate clines can be described by a pair of vectors, each with *n* slopes:

$$\boldsymbol{b_{N,t}} = \left(b_{N,1}, b_{N,2}, \dots, b_{N,n}\right) \qquad (S26a)$$

$$\boldsymbol{b_{I,t}} = \left(b_{I,1,t}, b_{I,2,t}, \dots, b_{I,n,t}\right) \qquad (S26b)$$

The "cosine similarity" between the vectors is:

$$\cos(\theta_t) = \frac{\sum_{i=1}^{n} b_{N,i} b_{I,i,t}}{\|\boldsymbol{b_N}\| \cdot \|\boldsymbol{b_{I,t}}\|} \qquad (S27)$$

where $\theta$ is the angle between the vectors, and $\|\boldsymbol{b_N}\| = \sqrt{\sum_{i=1}^{n} b_{N,i}^2}$ and $\|\boldsymbol{b_{I,t}}\| = \sqrt{\sum_{i=1}^{n} b_{I,i,t}^2}$ represent their magnitudes.

In this section, we focus on a special case of the model in which the traits are genetically independent of each other (thus, the single-trait cline models, presented above, predict multivariate clines), and introduced range trait clines are initially absent at time *t* = 0 ($b_0 = 0$ for each of the *n* traits). We later relax both assumptions.

##### Uniform population density model

Assuming uniform population density in each range, we substitute $b_{I,i,t} = B_{I,i}(1 - (1 - h_i^2\gamma_i)^t)$ and $b_{N,i} = B_{N,i}$ and the cosine similarity at time *t* becomes:

$$\cos(\theta_t) = \frac{\sum_{i=1}^{n} B_{N,i} B_{I,i}(1 - (1 - h_i^2\gamma_i)^t)}{\sqrt{\left(\sum_{i=1}^{n} B_{N,i}^2\right)\left(\sum_{i=1}^{n} B_{I,i}^2(1 - (1 - h_i^2\gamma_i)^t)^2\right)}} \qquad (S28)$$

In the limit of large *t*, we have:

$$\lim_{t\to\infty} \cos(\theta_t) = \cos\left(\theta_{eq.}\right) = \frac{\sum_{i=1}^{n} B_{N,i} B_{I,i}}{\sqrt{\left(\sum_{i=1}^{n} B_{N,i}^2\right)\left(\sum_{i=1}^{n} B_{I,i}^2\right)}} \qquad (S29)$$

In contrast, shortly after the introduction event, we can substitute $(1 - h_i^2\gamma_i)^t \approx 1 - th_i^2\gamma_i$ (valid when $th_i^2\gamma_i \ll 1$), which gives:

$$\lim_{t\to 0} \cos(\theta_t) \approx \frac{\sum_{i=1}^{n} B_{N,i}B_{I,i}h_i^2\gamma_i}{\sqrt{\left(\sum_{i=1}^{n} B_{N,i}^2\right)\left(\sum_{i=1}^{n} B_{I,i}^2(h_i^2\gamma_i)^2\right)}} \quad \text{(S30)}$$

Assuming the distribution of $h_i^2\gamma_i$ among traits is independent of $B_{N,i}$ and $B_{I,i}$, then we have:

$$\sum_{i=1}^{n} B_{N,i}B_{I,i}h_i^2\gamma_i = n\frac{1}{n}\sum_{i=1}^{n} B_{N,i}B_{I,i}h_i^2\gamma_i = n\,\mathrm{E}\left[B_{N,i}B_{I,i}\right]\mathrm{E}[h_i^2\gamma_i] = \mathrm{E}[h_i^2\gamma_i]\sum_{i=1}^{n} B_{N,i}B_{I,i} \quad \text{(S31)}$$

$$\sum_{i=1}^{n} B_{I,i}^2(h_i^2\gamma_i)^2 = n\frac{1}{n}\sum_{i=1}^{n} B_{I,i}^2(h_i^2\gamma_i)^2 = n\,\mathrm{E}\left[B_{I,i}^2\right]\mathrm{E}[(h_i^2\gamma_i)^2]$$

$$= \left(\mathrm{E}[h_i^2\gamma_i]^2 + \mathrm{var}[h_i^2\gamma_i]\right)\sum_{i=1}^{n} B_{I,i}^2 \quad \text{(S32)}$$

$$\lim_{t\to 0} \cos(\theta_t) \approx \frac{\mathrm{E}[h_i^2\gamma_i]}{\sqrt{\mathrm{E}[h_i^2\gamma_i]^2 + \mathrm{var}[h_i^2\gamma_i]}} \cdot \cos(\theta_{eq.}) = \frac{\cos(\theta_{eq.})}{\sqrt{1 + \frac{\mathrm{var}[h_i^2\gamma_i]}{\mathrm{E}[h_i^2\gamma_i]^2}}} \quad \text{(S33)}$$

A similar analysis applies for the entire range of *t*. Again, if we assume that the distribution of $h_i^2\gamma_i$ among traits is independent of $B_{N,i}$ and $B_{I,i}$, then:

$$\sum_{i=1}^{n} B_{N,i}B_{I,i}\left(1 - (1 - h_i^2\gamma_i)^t\right) = \frac{\sum_{i=1}^{n}\left(1 - (1 - h_i^2\gamma_i)^t\right)}{n}\sum_{i=1}^{n} B_{N,i}B_{I,i} \quad \text{(S34)}$$

$$\sum_{i=1}^{n} B_{I,i}^2\left(1 - (1 - h_i^2\gamma_i)^t\right)^2 = \frac{\sum_{i=1}^{n}\left(1 - (1 - h_i^2\gamma_i)^t\right)^2}{n}\sum_{i=1}^{n} B_{I,i}^2 \quad \text{(S35)}$$

$$\frac{\sum_{i=1}^{n}\left(1 - (1 - h_i^2\gamma_i)^t\right)}{n} = \mathrm{E}[1 - (1 - h_i^2\gamma_i)^t] \quad \text{(S36)}$$

$$\frac{\sum_{i=1}^{n}\left(1 - (1 - h_i^2\gamma_i)^t\right)^2}{n} = \mathrm{E}[1 - (1 - h_i^2\gamma_i)^t]^2\left(1 + \frac{\mathrm{var}[1 - (1 - h_i^2\gamma_i)^t]}{\mathrm{E}[1 - (1 - h_i^2\gamma_i)^t]^2}\right) \quad \text{(S37)}$$

$$\cos(\theta_t) = \frac{\cos(\theta_{eq.})}{\sqrt{1 + \frac{\mathrm{var}[1 - (1 - h_i^2\gamma_i)^t]}{\mathrm{E}[1 - (1 - h_i^2\gamma_i)^t]^2}}} \quad \text{(S38)}$$

which converges to the same limits as before:

$$\lim_{t\to 0}\cos(\theta_t) \approx \frac{\cos(\theta_{eq.})}{\sqrt{1+\frac{\mathrm{var}[h_i^2\gamma_i]}{\mathrm{E}[h_i^2\gamma_i]^2}}} \quad \text{(S39)}$$

$$\lim_{t\to\infty}\cos(\theta_t) = \cos(\theta_{eq.}) \quad \text{(S40)}$$

Thus, if there is substantial variation in $h_i^2\gamma_i$ among traits, we should see the $\cos(\theta)$ values for different species tend to increase with the time since the species introduction occurred (i.e., $\cos(\theta)$ should increase with *t*).

Assuming uniform population density in each range, then the relative magnitude of divergence in the introduced and native ranges is:

$$\frac{\|\boldsymbol{b_{I,t}}\|}{\|\boldsymbol{b_N}\|} = \frac{\sqrt{\sum_{i=1}^{n} b_{I,i,t}^2}}{\sqrt{\sum_{i=1}^{n} b_{N,i}^2}} = \frac{\sqrt{\sum_{i=1}^{n} B_{I,i}^2(1-(1-h_i^2\gamma_i)^t)^2}}{\sqrt{\sum_{i=1}^{n} B_{N,i}^2}} \quad \text{(S41)}$$

Assuming the distribution of $h_i^2\gamma_i$ among traits is independent of $B_{N,i}$ and $B_{I,i}$, then:

$$\sum_{i=1}^{n} B_{I,i}^2(1-(1-h_i^2\gamma_i)^t)^2 = \frac{\sum_{i=1}^{n}(1-(1-h_i^2\gamma_i)^t)^2}{n}\sum_{i=1}^{n} B_{I,i}^2$$

$$= \mathrm{E}[1-(1-h_i^2\gamma_i)^t]^2\left(1+\frac{\mathrm{var}[1-(1-h_i^2\gamma_i)^t]}{\mathrm{E}[1-(1-h_i^2\gamma_i)^t]^2}\right)\sum_{i=1}^{n} B_{I,i}^2 \quad \text{(S42)}$$

$$\frac{\|\boldsymbol{b_{I,t}}\|}{\|\boldsymbol{b_N}\|} = \frac{\|\boldsymbol{B_I}\|}{\|\boldsymbol{B_N}\|}\mathrm{E}[1-(1-h_i^2\gamma_i)^t]\sqrt{1+\frac{\mathrm{var}[1-(1-h_i^2\gamma_i)^t]}{\mathrm{E}[1-(1-h_i^2\gamma_i)^t]^2}} \quad \text{(S43)}$$

Immediately following the introduction, this simplifies approximately to:

$$\lim_{t\to 0}\frac{\|\boldsymbol{b_{I,t}}\|}{\|\boldsymbol{b_N}\|} \approx \frac{\|\boldsymbol{B_I}\|}{\|\boldsymbol{B_N}\|}t\,\mathrm{E}[h_i^2\gamma_i]\sqrt{1+\frac{\mathrm{var}[h_i^2\gamma_i]}{\mathrm{E}[h_i^2\gamma_i]^2}} \quad \text{(S44)}$$

which is valid as long as $(1-h_i^2\gamma_i)^t \approx 1-th_i^2\gamma_i$.

When there are initially no trait clines in the introduced range, as assumed in this section, then it becomes clear that the ratio of magnitudes of divergence is much more sensitive to $t$ than is the cosine similarity between orientations of divergence, and numerical results bear that out (see Fig. A1, below). If the product of heritability and strength of selection varies among traits, both $\cos(\theta_t)$ and $\frac{\|\boldsymbol{b}_{I,t}\|}{\|\boldsymbol{b}_N\|}$ are expected to increase over time. However, $\frac{\|\boldsymbol{b}_{I,t}\|}{\|\boldsymbol{b}_N\|}$ increases with $t$ regardless of the specific pattern of variability (or absence of variability) in $h_i^2\gamma_i$ across traits. Consequently, $\frac{\|\boldsymbol{b}_{I,t}\|}{\|\boldsymbol{b}_N\|}$ shows a much stronger increase over time than does $\cos(\theta_t)$, which is not consistent with our empirical finding that $\cos(\theta_t)$ increases substantially as a function of time since introduction, while $\frac{\|\boldsymbol{b}_{I,t}\|}{\|\boldsymbol{b}_N\|}$ is roughly constant.

**Abundant center population density model**

Assuming an abundant center population density distribution in each range, and that the native range populations are at equilibrium, then the cosine similarity at time $t$ becomes:

$$\cos(\theta_t) = \frac{\sum_{i=1}^{n} \frac{h_{N,i}^2\gamma_{N,i}B_{N,i}}{\sigma_N^2 + h_{N,i}^2\gamma_{N,i}} \frac{h_{I,i}^2\gamma_{I,i}B_{I,i}}{\sigma_I^2 + h_{I,i}^2\gamma_{I,i}} (1 - (1 - \sigma_I^2 - h_i^2\gamma_i)^t)}{\sqrt{\left(\sum_{i=1}^{n} \left(\frac{h_{N,i}^2\gamma_{N,i}B_{N,i}}{\sigma_N^2 + h_{N,i}^2\gamma_{N,i}}\right)^2\right)\left(\sum_{i=1}^{n} \left(\frac{h_{I,i}^2\gamma_{I,i}B_{I,i}}{\sigma_I^2 + h_{I,i}^2\gamma_{I,i}}\right)^2 (1 - (1 - \sigma_I^2 - h_i^2\gamma_i)^t)^2\right)}} \tag{S45}$$

and the ratio of magnitudes of clinal divergence is:

$$\frac{\|\boldsymbol{b}_{I,t}\|}{\|\boldsymbol{b}_N\|} = \frac{\sqrt{\sum_{i=1}^{n} \left(\frac{h_{I,i}^2\gamma_{I,i}B_{I,i}}{\sigma_I^2 + h_{I,i}^2\gamma_{I,i}}\right)^2 (1 - (1 - \sigma_I^2 - h_i^2\gamma_i)^t)^2}}{\sqrt{\sum_{i=1}^{n} \left(\frac{h_{N,i}^2\gamma_{N,i}B_{N,i}}{\sigma_N^2 + h_{N,i}^2\gamma_{N,i}}\right)^2}} \tag{S46}$$

At equilibrium, the cosine similarity and relative magnitudes simplify to:

$$\cos(\theta_{eq.}) = \frac{\sum_{i=1}^{n} \frac{h_{N,i}^2\gamma_{N,i}B_{N,i}}{\sigma_N^2 + h_{N,i}^2\gamma_{N,i}} \frac{h_{I,i}^2\gamma_{I,i}B_{I,i}}{\sigma_I^2 + h_{I,i}^2\gamma_{I,i}}}{\sqrt{\left(\sum_{i=1}^{n} \left(\frac{h_{N,i}^2\gamma_{N,i}B_{N,i}}{\sigma_N^2 + h_{N,i}^2\gamma_{N,i}}\right)^2\right)\left(\sum_{i=1}^{n} \left(\frac{h_{I,i}^2\gamma_{I,i}B_{I,i}}{\sigma_I^2 + h_{I,i}^2\gamma_{I,i}}\right)^2\right)}} \tag{S47}$$

$$\frac{\|\boldsymbol{b}_{\boldsymbol{I},\boldsymbol{eq.}}\|}{\|\boldsymbol{b}_{\boldsymbol{N}}\|}=\frac{\sqrt{\sum_{i=1}^{n}\left(\frac{h_{I,i}^2\gamma_{I,i}B_{I,i}}{\sigma_I^2+h_{I,i}^2\gamma_{I,i}}\right)^2}}{\sqrt{\sum_{i=1}^{n}\left(\frac{h_{N,i}^2\gamma_{N,i}B_{N,i}}{\sigma_N^2+h_{N,i}^2\gamma_{N,i}}\right)^2}} \tag{S48}$$

Assuming the distribution of $h_i^2\gamma_i$ among traits is independent of $B_{N,i}$ and $B_{I,i}$, and that the patterns of heritability, selection and dispersal do not differ between the ranges ($h_i^2 = h_{N,i}^2 = h_{I,i}^2$, $\gamma_i = \gamma_{N,i} = \gamma_{I,i}$, and $\sigma^2 = \sigma_N^2 = \sigma_I^2$), then we have:

$$\sum_{i=1}^{n}\left(\frac{h_i^2\gamma_i}{\sigma^2+h_i^2\gamma_i}\right)^2 B_{N,i}B_{I,i} = \mathrm{E}\left[\left(\frac{h_i^2\gamma_i}{\sigma^2+h_i^2\gamma_i}\right)^2\right]\sum_{i=1}^{n}B_{N,i}B_{I,i}$$

$$=\left(\mathrm{var}\left[\frac{h_i^2\gamma_i}{\sigma^2+h_i^2\gamma_i}\right]+\mathrm{E}\left[\frac{h_i^2\gamma_i}{\sigma^2+h_i^2\gamma_i}\right]^2\right)\sum_{i=1}^{n}B_{N,i}B_{I,i} \tag{S49}$$

$$\sum_{i=1}^{n}\left(\frac{h_i^2\gamma_i}{\sigma^2+h_i^2\gamma_i}\right)^2 B_{N,i}^2 = \left(\mathrm{var}\left[\frac{h_i^2\gamma_i}{\sigma^2+h_i^2\gamma_i}\right]+\mathrm{E}\left[\frac{h_i^2\gamma_i}{\sigma^2+h_i^2\gamma_i}\right]^2\right)\sum_{i=1}^{n}B_{N,i}^2 \tag{S50}$$

$$\sum_{i=1}^{n}\left(\frac{h_i^2\gamma_i}{\sigma^2+h_i^2\gamma_i}\right)^2 B_{I,i}^2 = \left(\mathrm{var}\left[\frac{h_i^2\gamma_i}{\sigma^2+h_i^2\gamma_i}\right]+\mathrm{E}\left[\frac{h_i^2\gamma_i}{\sigma^2+h_i^2\gamma_i}\right]^2\right)\sum_{i=1}^{n}B_{I,i}^2 \tag{S51}$$

and the equilibria simplify to those of the uniform density model:

$$\cos(\theta_{eq.}) = \frac{\sum_{i=1}^{n}B_{N,i}B_{I,i}}{\sqrt{\left(\sum_{i=1}^{n}B_{N,i}^2\right)\left(\sum_{i=1}^{n}B_{I,i}^2\right)}} \tag{S52}$$

$$\frac{\|\boldsymbol{b}_{\boldsymbol{I},\boldsymbol{eq.}}\|}{\|\boldsymbol{b}_{\boldsymbol{N}}\|}=\frac{\sqrt{\sum_{i=1}^{n}B_{I,i}^2}}{\sqrt{\sum_{i=1}^{n}B_{N,i}^2}} \tag{S53}$$

Numerical results show that the equilibria are reached slightly faster under the abundant center model (see Fig. A1), as also predicted in the univariate results (see Appendix 2).

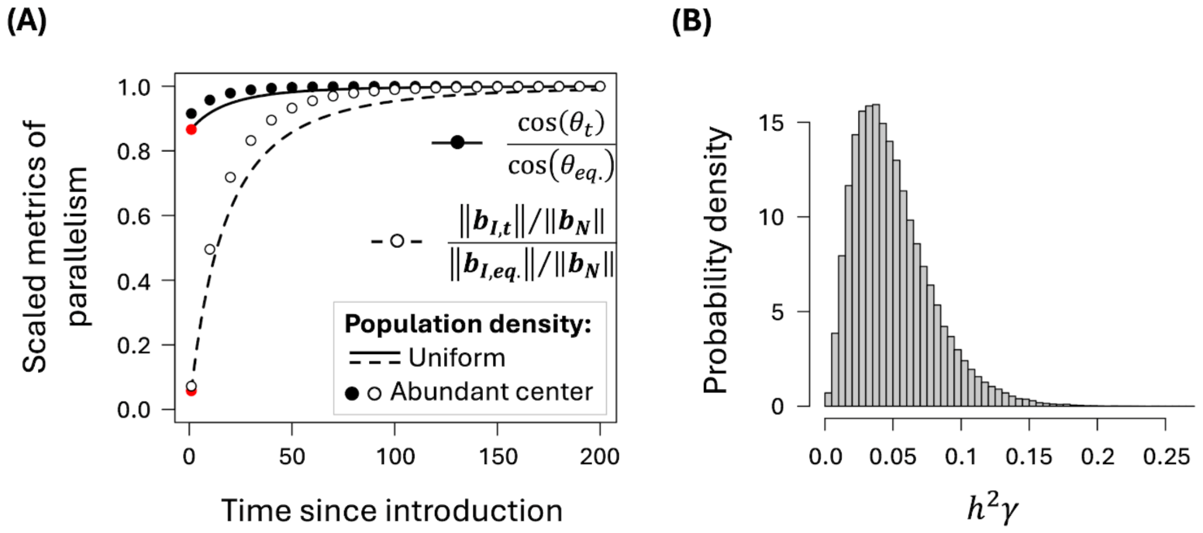


**Figure A1.** Changes in $\cos(\theta_t)$ and $\|\boldsymbol{b_{I,t}}\|/\|\boldsymbol{b_N}\|$ as a function of the time since introduction (left-hand panel), and the distribution of the product of heritability and the strength of selection across traits contributing to local adaptation (right-hand panel). Results assume that there are initially no introduced range trait clines at time $t$ = 0. In the left-hand panel, $B_{I,i}$ and $B_{N,i}$ follow a bivariate standard normal distribution with correlation coefficient of 0.5, the dispersal rate was fixed at $\sigma^2 =$ 0.01, and $h_i^2\gamma_i$ follow a gamma distribution with shape parameter $k$ = 3 and mean of $\mathrm{E}[h_i^2\gamma_i] = 0.05$. Parameters were randomly drawn for $10^5$ traits, and the trajectories of $\cos(\theta_t)$ and $\|\boldsymbol{b_{I,t}}\|/\|\boldsymbol{b_N}\|$ were calculated, with curves showing results for the uniform population density model, and black circles (filled and unfilled) showing results for the abundant center model. The red filled circles show analytical approximations for $\cos(\theta_t)$ and $\|\boldsymbol{b_{I,t}}\|/\|\boldsymbol{b_N}\|$ at generation $t$ = 1. For ease of comparison, results in the left-hand panel are scaled relative to the long-term equilibrium for each metric (hence, all results converge to one over time).

### Appendix 4: Serial bottlenecks and drift-induced clines

Here, we present a simple analytical approximation for the evolution of clines by drift during a range expansion. Our model is in the spirit of a simulation study by Colautti and Lau (2015), which assumed that a range expansion proceeds by sequential colonization and expansion events, each occurring in a stepwise manner across a linear array of adjacent habitat patches in the introduced range of a species. They also assumed that evolution of the trait proceeds by drift alone, as we do here as well. This model should therefore predict upper bounds for the initial frequency and steepness of counter-clines that arise during the range expansion. Counter-clines would be less frequent (and shallower in cases where they occur) if local selection were to play a substantial role in the initial formation of clines during the range expansion.

We assume that drift in a patch has a negligible effect on change in the trait mean once that patch has reached the local carrying capacity, but that drift can influence the trait mean before carrying capacity is reached. Suppose that an occupied patch, upon reaching carrying capacity, has a trait mean of $\bar{z}$. The additive genetic variance is $V_A$, which we assume is constant across populations. The population size dynamics during a colonization event of the next available adjacent patch are described by:

$$N_{e,t} = N_0 e^{rt} \tag{S71}$$

where $N_0$ is the founder population size for the next patch, $r$ is the rate of growth, and $t$ is the number of generations since the founder event initiating growth in the patch. Under this simple growth model, the local population grows exponentially until it reaches $K$, and it subsequently remains at $K$. The carrying capacity ($K$) is reached at generation $t = \frac{1}{r}\ln\left(\frac{K}{N_0}\right)$.

Following Lande (1976), evolutionary change in the trait mean during a single generation of drift is a normally distributed random variable with a mean of $\mathrm{E}[\Delta\bar{z}_t] = 0$ and a variance of $\mathrm{var}[\Delta\bar{z}_t] = V_A/N_e = h^2/N_e$ (the last expression applies because phenotypic variance is scaled to one in our models). The cumulative change during colonization of a single patch is therefore:

$$\Delta\bar{z}_{col} = \sum_{t=0}^{t=\frac{1}{r}\ln\left(\frac{K}{N_0}\right)} x_t \tag{S72}$$

where $x_t \sim N\left(\mathrm{E}[x_t] = 0; \mathrm{var}[x_t] = h^2/N_{e,t}\right)$. Consequently, the distribution of $\Delta\bar{z}_{col}$ will also be normally distributed with mean of zero and variance of:

$$\mathrm{var}[\Delta\bar{z}_{col}] = \sum_{t=0}^{t=\frac{1}{r}\ln\left(\frac{K}{N_0}\right)} \mathrm{var}[x_t] = \frac{h^2}{N_0} \sum_{t=0}^{t=\frac{1}{r}\ln\left(\frac{K}{N_0}\right)} e^{-rt} = \frac{h^2}{N_0} \cdot \frac{1 - e^{-\ln\left(\frac{K}{N_0}\right)-r}}{1 - e^{-r}} \tag{S73}$$

The last result implies a greater potential for drift during colonization when: (1) trait heritability is high; (2) population bottlenecks are severe; and (3) growth to carrying capacity is slow ($r$ is small and $K$ is high). A cline slope calculated from a pair of adjacent patches is:

$$\mathrm{slope} = \frac{y}{d} \tag{S74}$$

where $d$ is the distance between adjacent patches and $y$ is a normally distributed random variable: $y \sim N(0, \mathrm{var}[y] = \mathrm{var}[\Delta\bar{z}_{col}])$. If expansion across the introduced range proceeds by a series of colonization events across a linear array of equidistant habitat patches, then the variance of slopes between adjacent patches is equivalent to the velocity of a Brownian motion model of change over space. The velocity of the Brownian motion is:

$$v = \frac{1}{d^2}\mathrm{var}(y) = \frac{1}{d^2} \cdot \frac{h^2}{N_0} \cdot \frac{1 - e^{-\ln\left(\frac{K}{N_0}\right)-r}}{1 - e^{-r}} \tag{S75}$$

Once again, we see that greater rates of change in the trait mean across the introduced range are expected when bottlenecks are severe, growth in each habitat is slow, and trait heritability is high.

#### Appendix 5: Clinal divergence when the initial clines are drift-induced

We now revisit the multivariate model from Appendix 3, where traits are assumed to be genetically independent of each other. In this section we now allow for non-zero clines in the introduced range as an initial condition (i.e., at $t = 0$), and explore how metrices of parallelism change from the initial state to the equilibrium at migration-selection balance. As before, we assume that clines in the native range are at equilibrium, in which case the pattern of multivariate clinal divergence in the introduced relative to the native range is described by:

$$\frac{||\boldsymbol{b_I}||}{||\boldsymbol{b_N}||} = \frac{\sqrt{\sum_{j=1}^{n} b_{j,I}^2}}{\sqrt{\sum_{j=1}^{n} b_{j,N,eq.}^2}} \tag{S76}$$

and

$$\cos(\theta) = \frac{\sum_{j=1}^{n} b_{j,I} \cdot b_{j,N,eq.}}{\sqrt{\left(\sum_{j=1}^{n} b_{j,I}^2\right)\left(\sum_{j=1}^{n} b_{j,N,eq.}^2\right)}} \tag{S77}$$

With a uniform population density distribution following the range expansion, then using eq. (S6), the pattern of multivariate divergence is:

$$\frac{||\boldsymbol{b_I}||}{||\boldsymbol{b_N}||} = \frac{\sqrt{\sum_{j=1}^{n}\left(B_{j,I}\left(1-\left(1-h_j^2\gamma_j\right)^t\right)+b_{j,0}\left(1-h_j^2\gamma_j\right)^t\right)^2}}{\sqrt{\sum_{j=1}^{n} B_{j,N}^2}} \tag{S78}$$

and

$$\cos(\theta) = \frac{\sum_{j=1}^{n}\left(B_{j,I}B_{j,N}\left(1-\left(1-h_j^2\gamma_j\right)^t\right)+b_{j,0}B_{j,N}\left(1-h_j^2\gamma_j\right)^t\right)}{\sqrt{\left(\sum_{j=1}^{n}\left(B_{j,I}\left(1-\left(1-h_j^2\gamma_j\right)^t\right)+b_{j,0}\left(1-h_j^2\gamma_j\right)^t\right)^2\right)\left(\sum_{j=1}^{n} B_{j,N}^2\right)}} \tag{S79}$$

With selection sufficiently weak that $\left(1-h_j^2\gamma_j\right)^t \approx 1-h_j^2\gamma_j t$ then for small $t$, we have:

$$\lim_{t\to 0}\frac{||\boldsymbol{b_I}||}{||\boldsymbol{b_N}||} \approx \frac{\sqrt{\sum_{j=1}^{n} b_{j,0}^2}}{\sqrt{\sum_{j=1}^{n} B_{j,N}^2}} \tag{S80}$$

and

$$\lim_{t\to 0}\cos(\theta) \approx \frac{\sum_{j=1}^{n} b_{j,0}B_{j,N}}{\sqrt{\left(\sum_{j=1}^{n} b_{j,0}^2\right)\left(\sum_{j=1}^{n} B_{j,N}^2\right)}} \tag{S81}$$

When initial clines are caused by drift during the range expansion, then the values of $b_{j,0}$ will be independently drawn from a normal distribution with a mean of zero and variance, which we label here as $V_b$, whose magnitude depends on the severity of drift during the range expansion (see Appendix 5). If the equilibrium cline slopes in the native range are drawn from a normal distribution with mean of zero and variance of $V_B$, then for large $n$, we have:

$$\lim_{t\to 0}\frac{||\boldsymbol{b_I}||}{||\boldsymbol{b_N}||} \approx \lim_{t\to 0}\mathrm{E}\left[\frac{||\boldsymbol{b_I}||}{||\boldsymbol{b_N}||}\right] \approx \frac{\sqrt{V_b}}{\sqrt{V_B}} \tag{S82}$$

and

$$\lim_{t\to 0}\cos(\theta) \approx \lim_{t\to 0}\mathrm{E}[\cos(\theta)] = 0 \tag{S83}$$

The prediction for $\cos(\theta)$ follows from the fact that $b_{j,0}$ and $B_{j,N}$ are uncorrelated random variables. The prediction for the ratio of magnitudes is obtained as follows. With $b_{j,0}$ and $B_{j,N}$ each normally distributed with means of zero and variances of $V_b$ and $V_B$, respectively, then we have:

$$\lim_{t\to 0}\frac{||\boldsymbol{b_I}||}{||\boldsymbol{b_N}||} \approx \frac{\sqrt{\sum_{j=1}^{n} b_{j,0}^2}}{\sqrt{\sum_{j=1}^{n} B_{j,N}^2}} = \frac{\sqrt{V_b}}{\sqrt{V_B}}\cdot\frac{X}{Y} \tag{S84}$$

where $X = \sqrt{\sum_{j=1}^{n}\frac{b_{j,0}^2}{V_b}}$ and $Y = \sqrt{\sum_{j=1}^{n}\frac{B_{j,N}^2}{V_B}}$ are independent chi-distributed random variables with $n$ degrees of freedom. The expected value of the ratio can then be approximated using the Taylor series expansion:

$$\lim_{t\to 0}\mathrm{E}\left[\frac{||\boldsymbol{b_I}||}{||\boldsymbol{b_N}||}\right] \approx \frac{\sqrt{V_b}}{\sqrt{V_B}}\mathrm{E}\left[\frac{\sqrt{\sum_{j=1}^{n}\frac{b_{j,0}^2}{V_b}}}{\sqrt{\sum_{j=1}^{n}\frac{B_{j,N}^2}{V_B}}}\right] \approx \frac{\sqrt{V_b}}{\sqrt{V_B}}\left(1+\frac{\mathrm{var}[Y]}{\mathrm{E}[Y]^2}\right) \approx \frac{\sqrt{V_b}}{\sqrt{V_B}} \tag{S85}$$

The final approximation, appropriate when $n$ is large, yields eq. (S82).

Overall, we see that the ratio of magnitudes of clinal divergence in the introduced relative to the native range should initially be high relative to $\cos(\theta)$ if drift contributes substantially to cline formation during the range expansion. The long-run change in the relative magnitudes may ultimately conform between ranges (i.e., $||\boldsymbol{b_I}||$ may evolve to match $||\boldsymbol{b_N}||$) if similar cline magnitudes are favoured in the two ranges. By driving up the initial value of the ratio of magnitudes, drift-induced clines can result in $||\boldsymbol{b_I}||/||\boldsymbol{b_N}||$ changing very little over time. Drift has the opposite effect on $\cos(\theta)$, which may change from $\cos(\theta) \approx 0$ to $\cos(\theta) \approx 1$ over time if selection favours parallel cline patterns between the ranges.

#### Appendix 6: Effects of genetic correlations between traits

To explore the effects of genetic correlations between traits on the evolution of trait clines, we follow the general multivariate approach outlined in Duputié et al. (2012). The evolutionary dynamics for a pair of traits (with 1 and 2 subscripts referring to "trait 1" and "trait 2") are described by the set of difference equations:

$$\Delta\bar{z}_1(x,t) = \frac{\sigma^2}{2}\frac{\partial^2\bar{z}_1(x,t)}{\partial x^2} + \sigma^2\frac{\partial\log\big(N(x)\big)}{\partial x}\frac{\partial\bar{z}_1(x,t)}{\partial x} + G_1\beta_1(x,t) + G_{12}\beta_2(x,t) \quad \text{(S54)}$$

$$\Delta\bar{z}_2(x,t) = \frac{\sigma^2}{2}\frac{\partial^2\bar{z}_2(x,t)}{\partial x^2} + \sigma^2\frac{\partial\log\big(N(x)\big)}{\partial x}\frac{\partial\bar{z}_2(x,t)}{\partial x} + G_2\beta_2(x,t) + G_{12}\beta_1(x,t) \quad \text{(S55)}$$

where $\beta_1(x,t)$ and $\beta_2(x,t)$ are the selection gradients at location $x$ in the range for traits 1 and 2, $G_1$ and $G_2$ are the additive genetic variances for the traits, and $G_{12}$ is the genetic covariance between the traits. Assuming, for simplicity and consistency with the models presented above, we set the trait variance to unity ($V_P = 1$ for each trait), in which case the additive genetic variances are equivalent to heritabilities for the pair of traits ($h_1^2 = G_1$, $h_2^2 = G_2$), and the covariance is a function of the genetic correlation between traits ($\rho$) and the trait-specific heritabilities: $G_{12} = \rho\sqrt{h_1^2 h_2^2}$. With linear shifts in the trait optima across the range, the selection gradients for each trait are:

$$\beta_1(x,t) = \gamma_1\big(B_1 x - \bar{z}_1(x,t)\big) \quad \text{(S56)}$$

$$\beta_2(x,t) = \gamma_2\big(B_2 x - \bar{z}_2(x,t)\big) \quad \text{(S57)}$$

where $\gamma_1$ and $\gamma_2$ denote the strengths of stabilizing selection for traits 1 and 2, respectively. Under a uniform density model with weak range edge effects, and thus nearly linear clines across the range, we can approximate the dynamics as follows:

$$\Delta\bar{z}_1(x,t) = h_1^2\gamma_1\big(B_1 x - \bar{z}_1(x,t)\big) + \gamma_2\big(B_2 x - \bar{z}_2(x,t)\big)\rho\sqrt{h_1^2 h_2^2} \quad \text{(S58)}$$

$$\Delta\bar{z}_2(x,t) = h_2^2\gamma_2\big(B_2 x - \bar{z}_2(x,t)\big) + \gamma_1\big(B_1 x - \bar{z}_1(x,t)\big)\rho\sqrt{h_1^2 h_2^2} \quad \text{(S59)}$$

To facilitate the analysis, we assume that the heritability and strength of stabilizing selection is the same between traits, but the slopes of their gradients differ ($h^2 = h_1^2 = h_2^2$; $\gamma = \gamma_1 = \gamma_2$; $B_1 \neq B_2$), in which case we can write:

$$\Delta\bar{z}_1(x,t) = h^2\gamma\big(B_1 x - \bar{z}_1(x,t)\big) + h^2\gamma\big(B_2 x - \bar{z}_2(x,t)\big)\rho \tag{S60}$$

$$\Delta\bar{z}_2(x,t) = h^2\gamma\big(B_2 x - \bar{z}_2(x,t)\big) + h^2\gamma\big(B_1 x - \bar{z}_1(x,t)\big)\rho \tag{S61}$$

Defining the variables $\bar{z}_{avg}(x,t) = \big(\bar{z}_1(x,t) + \bar{z}_2(x,t)\big)/2$ and $\bar{z}_{diff}(x,t) = \bar{z}_1(x,t) - \bar{z}_2(x,t)$, we can write difference equations that describe changes in the trait averages and trait differences, which are decoupled from one another under the stated assumptions:

$$\Delta\bar{z}_{avg}(x,t) = \frac{1}{2}\big(\Delta\bar{z}_1(x,t) + \Delta\bar{z}_2(x,t)\big) = h^2\gamma\left(\frac{1}{2}(B_1 + B_2)x - \bar{z}_{avg}(x,t)\right)(1+\rho) \tag{S62}$$

$$\Delta\bar{z}_{diff}(x,t) = \Delta\bar{z}_1(x,t) - \Delta\bar{z}_2(x,t) = h^2\gamma\left((B_1 - B_2)x - \bar{z}_{diff}(x,t)\right)(1-\rho) \tag{S63}$$

The general solutions to the difference equations are:

$$\bar{z}_{avg}(x,t) = \frac{1}{2}(B_1 + B_2)x - \left(\frac{1}{2}(B_1 + B_2)x - \bar{z}_{avg}(x,0)\right)\left(1 - h^2\gamma(1+\rho)\right)^t \tag{S64}$$

$$\bar{z}_{diff}(x,t) = (B_1 - B_2)x - \left((B_1 - B_2)x - \bar{z}_{diff}(x,0)\right)\left(1 - h^2\gamma(1-\rho)\right)^t \tag{S65}$$

Assuming that there is initially no cline for either trait at the time of the introduction ($\bar{z}_{avg}(x,0) = \bar{z}_{diff}(x,0) = 0$), then we can write the general solutions for the individual trait clines as:

$$\bar{z}_1(x,t) = \bar{z}_{avg}(x,t) + \frac{1}{2}\bar{z}_{diff}(x,t)$$

$$= B_1 x - \frac{1}{2}(B_1 - B_2)x\big(1 - h^2\gamma(1-\rho)\big)^t - \frac{1}{2}(B_1 + B_2)x\big(1 - h^2\gamma(1+\rho)\big)^t \tag{S66}$$

$$\bar{z}_2(x,t) = \bar{z}_{avg}(x,t) - \frac{1}{2}\bar{z}_{diff}(x,t)$$

$$= B_2 x + \frac{1}{2}(B_1 - B_2)x\big(1 - h^2\gamma(1-\rho)\big)^t - \frac{1}{2}(B_1 + B_2)x\big(1 - h^2\gamma(1+\rho)\big)^t \tag{S67}$$

Note that in the absence of a genetic correlation between the traits ($\rho = 0$), the solutions simplify to $\bar{z}_1(x,t) = B_1 x(1 - (1 - h^2\gamma)^t)$ and $\bar{z}_2(x,t) = B_2 x(1 - (1 - h^2\gamma)^t)$, which matches our earlier univariate results. The cline slopes are:

$$b_{1,t} = \frac{d\bar{z}_1(x,t)}{dx}$$

$$= B_1 - \frac{1}{2}(B_1 - B_2)\big(1 - h^2\gamma(1-\rho)\big)^t - \frac{1}{2}(B_1 + B_2)\big(1 - h^2\gamma(1+\rho)\big)^t \qquad \text{(S68)}$$

$$b_{2,t} = \frac{d\bar{z}_2(x,t)}{dx}$$

$$= B_2 + \frac{1}{2}(B_1 - B_2)\big(1 - h^2\gamma(1-\rho)\big)^t - \frac{1}{2}(B_1 + B_2)\big(1 - h^2\gamma(1+\rho)\big)^t \qquad \text{(S69)}$$

Assuming that genetic constraints to the evolution of the trait pair are not absolute (*i.e.*, the genetic correlation is not perfect; $|\rho| < 1$), then each trait will eventually evolve to an equilibrium where its cline closely tracks the optimum for the trait ($\bar{z}_1^*(x,t) = B_1 x$ and $\bar{z}_2^*(x,t) = B_2 x$ at equilibrium). Although both traits closely track their optima at equilibrium, it is possible for counter-gradient clines to evolve—*i.e.*, trait clines that are in the opposite direction of the change in the optimum for the trait—during the initial approach of an introduced population to the equilibrium in the new range. For example, consider the case where the pair of traits has a positive genetic correlation ($\rho > 0$) and selection favours clines with opposite slopes for the pair of traits (let $B_1 > 0$ and $B_2 < 0$, with $b_1/b_2 = B_1/B_2$ representing the ratio). If $|B_1| > |B_2|$ and $h^2\gamma \ll 1$, then the first trait will always evolve towards its equilibrium. A counter-gradient will initially evolve for the second trait under the condition:

$$\rho > -\frac{B_2}{B_1} \qquad \text{(S70)}$$

Thus, counter-clines can occur shortly after an introduction if the genetic correlation between the traits is sufficiently strong and environmental gradients have opposite signs but different magnitudes. The same can occur when the genetic correlation is strongly negative, and the environmental gradients have the same sign but different magnitudes. Predictions of the model are shown in Figure A2 (immediately below), where strong selection for local adaptation in one trait (trait 1, which has a cline slope of $b_{I,1}$ and $b_{N,1}$ in the introduced and native ranges, respectively) leads to a counter-cline in a second trait that is genetically correlated with the first (the second trait, trait 2, has a cline slope of $b_{I,2}$ and $b_{N,2}$ in the introduced and native ranges).

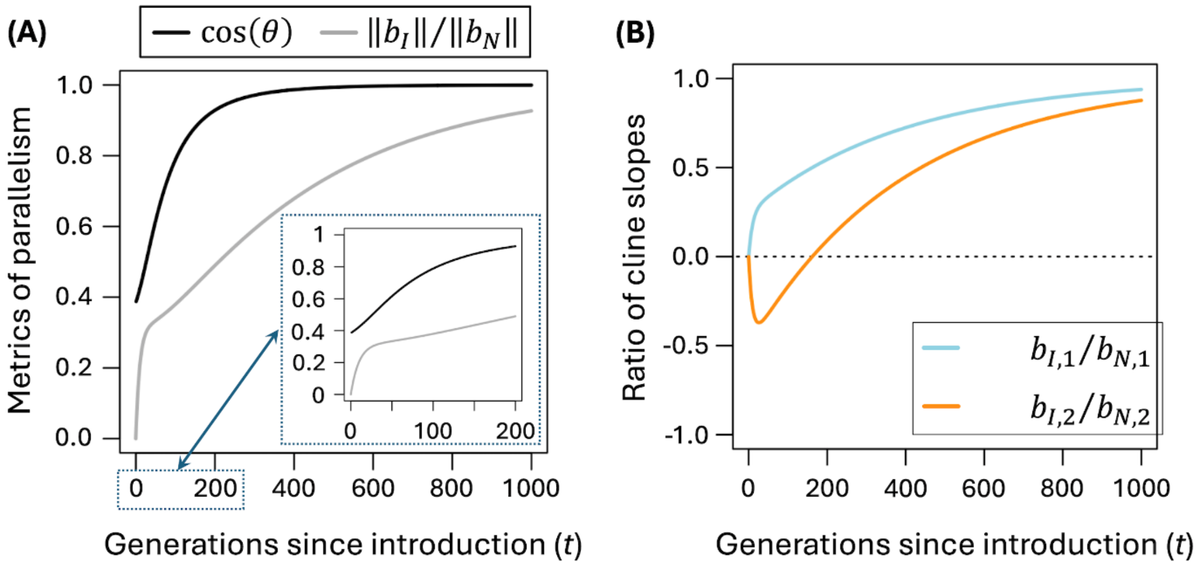


**Figure A2.** Evolution of clines for a pair of genetically correlated traits. The plots show a scenario in which the optima for the pair of traits shift in opposite directions, with $B_1 = 2$ and $B_1 = -1$, and the traits are otherwise subject to the same strength of stabilizing selection ($\gamma = 0.05$). Heritability for each trait is $h^2 = 1$ and the genetic correlation between the traits is strongly positive ($\rho = 0.95$). Parameters of selection and variation are assumed to be identical between the native and introduced ranges, so that perfect parallelism ($||\boldsymbol{b_I}||/||\boldsymbol{b_N}||$ and $\cos(\theta) = 1$) is predicted at equilibrium. Panel **(A)** shows the evolution of $\cos(\theta)$ and $||\boldsymbol{b_I}||/||\boldsymbol{b_N}||$ across 1000 generations since the introduction at generation $t = 0$, with the inset highlighting the predictions for the first 200 generations. Panel **(B)** shows the evolution of clines for each trait—including a counter-cline that arises early for trait 2 but ultimately resolves as introduced populations approach migration-selection equilibrium.

#### Supplementary Figures

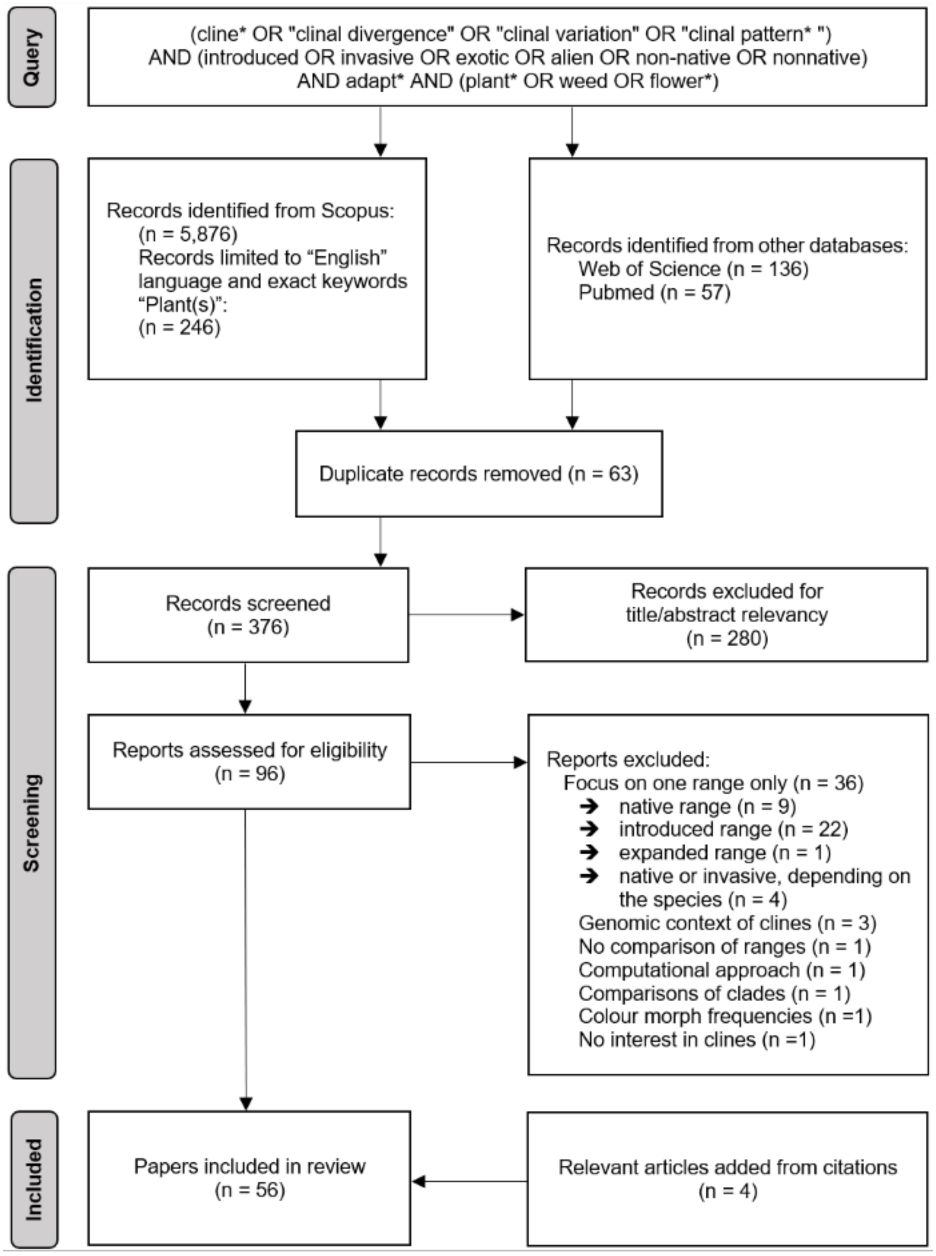


**Figure S1.** A PRISMA (Preferred Reporting Items for Systematic Reviews and Meta-Analyses) flow diagram of the studies comparing native and introduced clines in plant species.

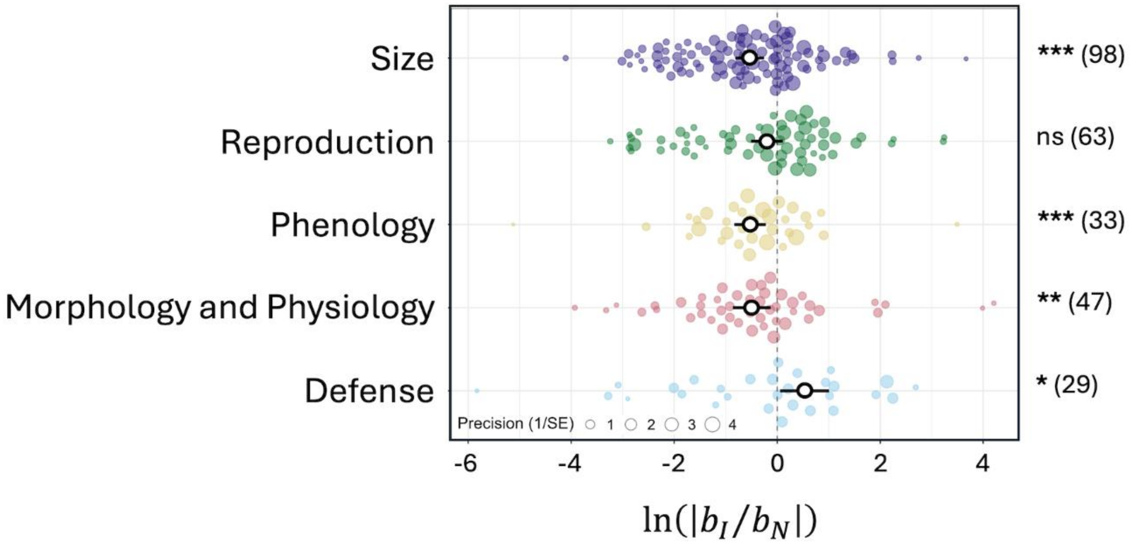


**Figure S2**. Ratios of absolute values of clines slopes in the introduced versus native range. Analyses are based on traits with a statistically significant cline in at least one range (as outlined in Fig. 3 of the main text), which includes cases where $b_I/b_N > 0$ (as in Fig. 4B-C of the main text) and where $b_I/b_N < 0$. Individual effect sizes (coloured circles) for the five trait categories are plotted with circle size denoting precision (inverse variance). Open circles denote mean effect sizes and whiskers are the 95% confidence intervals. The number of traits in each category is shown in brackets, with asterisks indicating significant deviations from $\ln(|b_I/b_N|) = 0$ (*$P < 0.05$, **$P < 0.01$, ***$P < 0.001$).

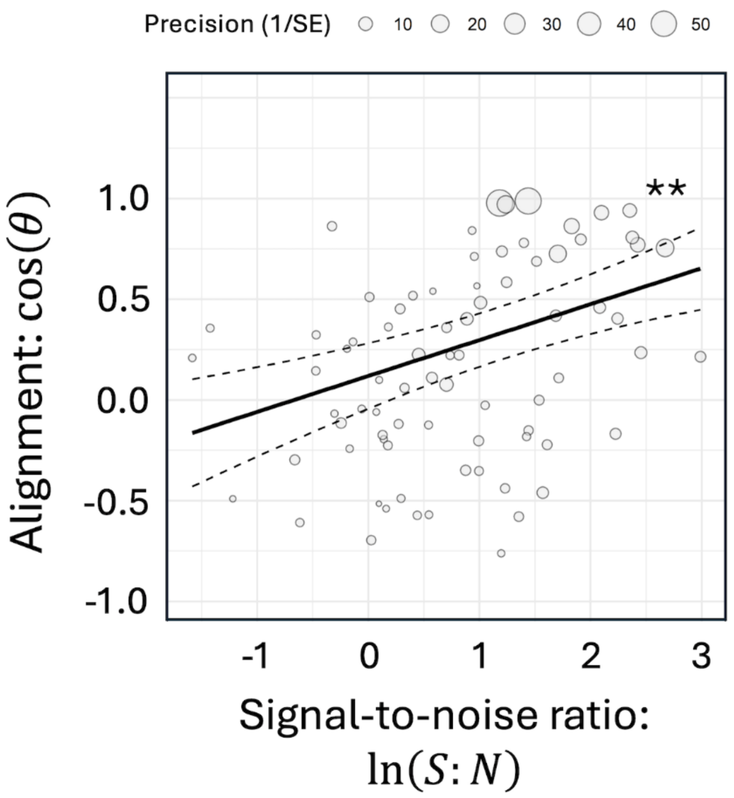


**Figure S3**: Relationship between vector alignment ($\cos\theta$) and signal-to-noise ratio ($\ln(S{:}N)$, with meta-analytic data shown as bubble plots. Asterisks indicate a significant association with time (** $P$ < 0.001).

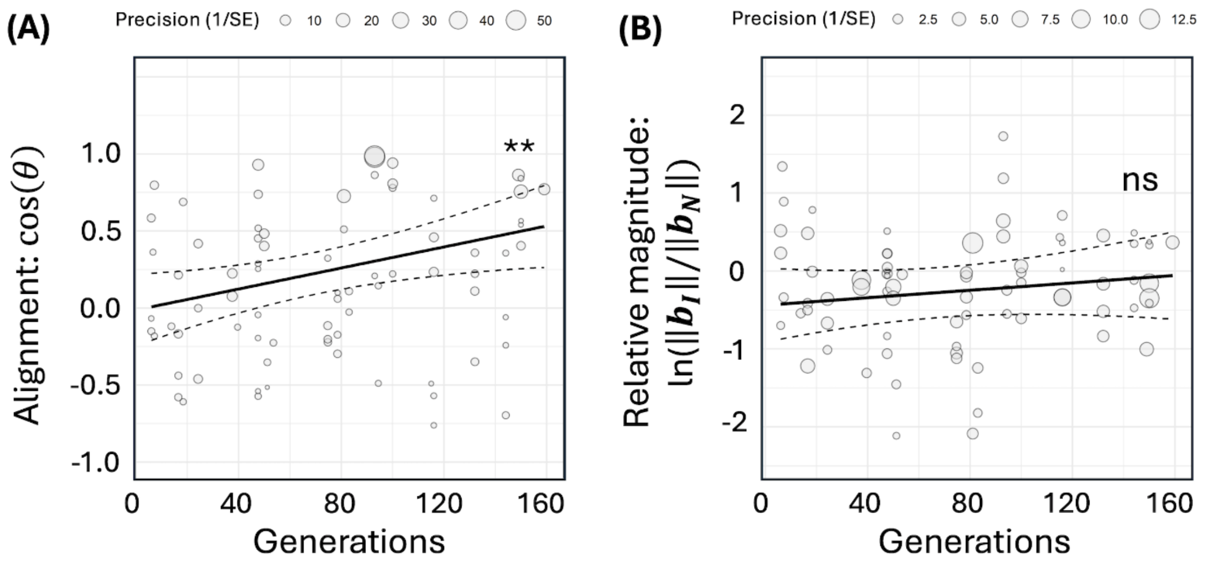


**Figure S4**: Associations between the number of generations since introduction and **(A)** cos($\theta$) or **(B)** ln($\|\boldsymbol{b_I}\|/\|\boldsymbol{b_N}\|$) with an outlier (generations > 200) removed. Meta-analytic data are shown as bubble plots. Asterisks indicate a significant association with time (** $P < 0.001$).

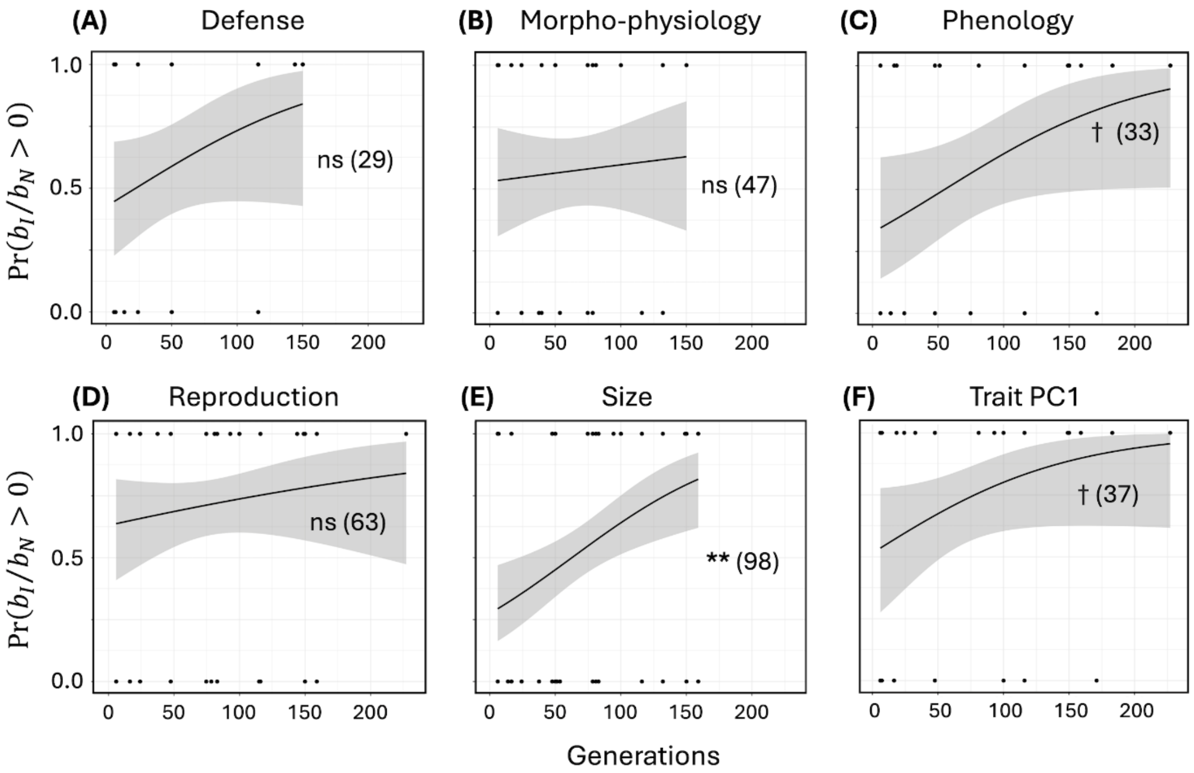


**Figure S5**. Generalized linear model of the probability of having $b_I/b_N > 0$, for each trait category and trait PC1, as a function of the number of generations since introduction ($b_I$ and $b_N$ are estimates of the cline slopes in the introduced and native ranges, respectively). The number of effect sizes is given in brackets. Symbols indicate significance: † $P < 0.1$, ** $P < 0.01$.

**Box S1.** List of the 34 references that met our inclusion criteria for the meta-analysis.

#### Supplementary Tables

**Table S1.** Dataset of the meta-analysis. The references of the paper IDs can be found in Box S1. When two dates appear, it means the paper focuses on introduced ranges with different times of introduction. "Generation number" refers to the number of generations since the introduction.

| Paper ID | Family | Species | Introduction year | Generations number |
|---|---|---|---|---|
| 1 | Asteraceae | *Lactuca serriola* | 1883 | 50 |
| 2 | Plantaginaceae | *Plantago lanceolata* | 1743 | 75.5 |
| 3 | Asteraceae | *Lactuca serriola* | 1770 | 95.6 |
| 4 | Brassicaceae | *Brassica tournefortii* | 1927 | 92 |
| 5 | Asteraceae | *Senecio pterophorus* | 1900/1982 | 25.8/9.3 |
| 6 | Poaceae | *Spartina alterniflora* | 1979 | 20.7 |
| 7 | Asteraceae | *Senecio pterophorus* | 1900/1982 | 25.6/7.3 |
| 8 | Scrophulariaceae | *Verbascum thapsus* | 1600 | 146.2 |
| 9 | Asteraceae | *Solidago altissima* | 1900 | 7 |
| 10 | Poaceae | *Bromus tectorum* | 1860 | 164 |
| 11 | Poaceae | *Phragmites australis* | 1828 | 43.6 |
| 12 | Fabaceae | *Medicago polymorpha* | 1832 | 186 |
| 13 | Fabaceae | *Trifolium repens* | 1665 | 34.3 |
| 14 | Asteraceae | *Helianthus annuus* | 1922 | 97 |
| 15 | Asteraceae | *Cirsium arvense* | 1822 | 83.6 |
| 16 | Asteraceae | *Chromolaena odorata* | 1934 | 16 |
| 17, 18 | Poaceae | *Spartina alterniflora* | 1979 | 19.3 |
| 19 | Poaceae | *Spartina alterniflora* | 1979 | 20.3 |
| 20 | Hypericaceae | *Hypericum perforatum* | 1748 | 102.4 |
| 21 | Asteraceae | *Ambrosia artemisiifolia* | 1863 | 157 |
| 22 | Plantaginaceae | *Plantago lanceolata* | 1839 | 52 |

| | | | | |
|---|---|---|---|---|
| 23 | Fabaceae | *Lotus corniculatus* | 1901 | 56 |
| 24 | Asteraceae | *Conyza canadensis* | 1700 | 227.1 |
| 25 | Brassicaceae | *Arabidopsis thaliana* | 1837 | 175 |
| 26 | Asteraceae | *Solidago altissima* | 1900 | 7.5 |
| 27 | Asteraceae | *Ambrosia artemisiifolia* | 1930 | 89 |
| 28 | Asteraceae | *Centaurea diffusa* | 1907 | 54.2 |
| 29 | Asteraceae | *Centaurea diffusa* | 1907 | 54.7 |
| 30, 31 | Asteraceae | *Ambrosia artemisiifolia* | 1863/1897 | 156/122 |
| 32 | Phrymaceae | *Mimulus guttatus* | 1824 | 38.8 |
| 33 | Polygonaceae | *Polygonum cespitosum* | 1927 | 95 |
| 34 | Amaranthaceae | *Alternanthera philoxeroides* | 1930 | 136.5 |
| 35 | Asteraceae | *Senecio vulgaris* | 1890 | 131 |

**Table S2.** Traits measured by the 34 papers that met the inclusion criteria, with the corresponding trait categories.

| **Trait measured** | **Trait category** |
|---|---|
| Abscisic acid | Defence |
| Aucubin | Defence |
| Carbon | Defence |
| Carbon/Nitrogen ratio | Defence |
| Catalpol | Defence |
| Cellulose | Defence |
| Diversity of pyrrolizidine alkaloids in seed | Defence |
| Diversity of volatile organic compounds | Defence |
| Flavonoids | Defence |
| Herbivory resistance (number of herbivores) | Defence |
| Jasmonate | Defence |
| Leaf area lost | Defence |
| Leaf consumed | Defence |
| Leaf chewing damage | Defence |
| Leaf thickness | Defence |
| Nitrogen | Defence |
| Phenolic concentration | Defence |
| Phenolic richness | Defence |
| Pyrrolizidine alkaloids in leaf (relative abundance) | Defence |
| Total phenolic content | Defence |
| Total pyrrolizidine alkaloids in leaf | Defence |
| Total volatile organic compounds | Defence |
| Trichome.density | Defence |
| Triterpenoidsaponins | Defence |
| Verbascoside | Defence |
| Aboveground biomass/Belowground biomass ratio | Morpho-physiology |
| Anther-stigma separation | Morpho-physiology |

| | |
|---|---|
| Branch architecture | Morpho-physiology |
| Branch/stem ratio | Morpho-physiology |
| Corolla length/Corolla width ratio | Morpho-physiology |
| Corolla width | Morpho-physiology |
| delta-13-C (ratio of carbon isotopes) | Morpho-physiology |
| Leaf area | Morpho-physiology |
| Leaf cost/Carbon accumulation ratio | Morpho-physiology |
| Leaf length | Morpho-physiology |
| Leaf mass area | Morpho-physiology |
| Leaf shape | Morpho-physiology |
| Leaf traits | Morpho-physiology |
| Leaf width | Morpho-physiology |
| Longest leaf | Morpho-physiology |
| Root-to-shoot ratio | Morpho-physiology |
| Soil moisture at wilt | Morpho-physiology |
| Specific leaf area | Morpho-physiology |
| Specific stem length | Morpho-physiology |
| Stomatal guard cell size | Morpho-physiology |
| Anthesis time | Phenology |
| Bolt time | Phenology |
| Bolt to bud | Phenology |
| Bud time | Phenology |
| Bud to flower | Phenology |
| Dichogamy | Phenology |
| Flower time | Phenology |
| Seed time | Phenology |
| Senescence time | Phenology |
| Sown to bolt | Phenology |
| Timson index | Phenology |
| Withered shoots percentage | Phenology |

| | |
|---|---|
| Awn length | Reproduction |
| Capitula number | Reproduction |
| Capsule number | Reproduction |
| Flower biomass | Reproduction |
| Flower density | Reproduction |
| Flower number | Reproduction |
| Flowering period | Reproduction |
| Flowered or not | Reproduction |
| Floral sex allocation | Reproduction |
| Fruit number | Reproduction |
| Inflorescence length | Reproduction |
| Proportion of germinated seeds | Reproduction |
| Relative reproductive biomass | Reproduction |
| Reproductive allocation | Reproduction |
| Reproductive biomass | Reproduction |
| Seed head number | Reproduction |
| Seed length | Reproduction |
| Seed mass | Reproduction |
| Seed number | Reproduction |
| Seed set percentage | Reproduction |
| Seed width | Reproduction |
| Survival | Reproduction |
| Total reproductive biomass | Reproduction |
| Aboveground biomass | Size |
| Adult-seedling height difference | Size |
| Belowground biomass | Size |
| Branch number | Size |
| Canopy area | Size |
| Elongation rate | Size |
| Height | Size |

| | |
|---|---|
| Leaf biomass | Size |
| Leaf dry matter content | Size |
| Leaf number | Size |
| Leaf water content | Size |
| Longest shoot | Size |
| Maximum growth rate | Size |
| Panicle height | Size |
| Photosynthetic rate | Size |
| Plant volume | Size |
| Root crown diameter | Size |
| Rosette area | Size |
| Rosette maximum diameter | Size |
| Rosette minimum diameter | Size |
| Shoot number | Size |
| Sparseness | Size |
| Spread area | Size |
| Stem biomass | Size |
| Stem diameter | Size |
| Stolon length | Size |
| Total biomass | Size |

**Table S3.** Overall effect size of multivariate and univariate parameters across all samples. A significant QE indicates that the residual heterogeneity can be explained by factors not included in the model. $I^2$ is the total heterogeneity at each random effect level, $\mathbf{b_I}$ and $\mathbf{b_N}$ are vectors of introduced and native cline slope estimates, cos(θ) represents the correlation of cline slopes between the ranges (θ being the angle between vectors), and $\ln(\|\mathbf{b_I}\|/\|\mathbf{b_N}\|)$ shows the log ratio of the cline vector magnitudes. Values, $b_I$ and $b_N$, are univariate cline slope estimates in the introduced and native ranges, respectively. CI = confidence interval. Asterisks indicate significance: *** $P$ <0.001.

| | Pooled effect size | 95% CI | $I^2_{\text{level 2}}$ (%) | $I^2_{\text{level 3}}$ (%) | QE |
|---|---|---|---|---|---|
| **Multivariate analysis** | | | | | |
| $\cos(\theta)$ | 0.30 | [0.13, 0.46] | 94.8 | 0 | 842 *** |
| $\ln(\|\mathbf{b_I}\|/\|\mathbf{b_N}\|)$ | -0.30 | [-0.58, -0.02] | 91 | 0 | 374*** |
| **Univariate analysis** | | | | | |
| $\ln(b_I/b_N)$ | -0.22 | [-0.47, 0.02] | 2.12 | 35.98 | 233*** |

**Table S4.** Variation in multivariate vector alignment ($\cos(\theta)$) was tested against six moderators, including: number of generations since introduction ('generations'), categories of environmental gradient used by authors ('gradient category'), self-compatibility ('mating system'), whether species were introduced across northern and southern hemispheres ('hemisphere'), proportion of environmental gradient shared between native and introduced ranges ('gradient overlap'), the number of traits examined ('trait number'), and log-transformed signal-to-noise ratio ('log-S:N'). . A model selection with *glmulti* based on the second order Akaike information criterion (AICc) found the best model to include generations, gradient overlap, and log-S:N when the analysis included all samples (**A**: model fit of AICc = 99.0, $F_{3,74}$ = 11.0, P < 0.0001, QE = 706.1), and when an outlier sample was excluded (**B**: AICc = 98.3, $F_{3,73}$ = 10.0, P < 0.0001, QE = 700.7). Tables **C** & **D** show results of full models when analysis included all samples (**C**: model fit: AICc = 111.5, $F_{10,67}$ = 3.2, P < 0.0001, QE = 256.3), and when analysis excluded an outlier (**D**: AICc = 109.3, $F_{10,66}$ = 3.1, P < 0.0001, QE = 255.9). Bold letters indicate statistical significance.

##### *A: Best model with all samples*

| | *Estimate* | *SE* | *t-value* | *df* | *P-value* |
|---|---|---|---|---|---|
| **Generations** | **0.0040** | **0.0012** | **3.49** | 74 | **0.0008** |
| Gradient overlap | -0.48 | 0.29 | -1.65 | 74 | 0.10 |
| **Log-S:N** | **0.18** | **0.042** | **4.25** | 74 | **< 0.0001** |

##### *B: Best model excluding an outlier*

| | *Estimate* | *SE* | *t-value* | *df* | *P-value* |
|---|---|---|---|---|---|
| **Generations** | **0.0034** | **0.0013** | **2.66** | 73 | **0.009** |
| Gradient overlap | -0.57 | 0.30 | -1.86 | 73 | 0.07 |
| **Log-S:N** | **0.18** | **0.042** | **4.31** | 73 | **< 0.0001** |

##### *C: Full model with all samples included*

| | *Estimate* | *SE* | *t-value* | *df* | *P-value* |
|---|---|---|---|---|---|
| **Generations** | **0.0048** | **0.0016** | **2.98** | 67 | **0.0041** |
| Gradient category (aridity) | 0.026 | 0.24 | 0.11 | 67 | 0.91 |
| Gradient category (BioClim) | -0.29 | 0.29 | -0.98 | 67 | 0.33 |
| Gradient category (elevation) | -0.11 | 0.44 | -0.25 | 67 | 0.80 |
| Gradient category (temperature) | 0.14 | 0.25 | 0.56 | 67 | 0.58 |
| Mating system | 0.0064 | 0.089 | 0.072 | 67 | 0.94 |
| Hemisphere | 0.088 | 0.11 | 0.78 | 67 | 0.44 |
| Gradient overlap | -0.60 | 0.43 | -1.39 | 67 | 0.17 |
| Trait number | -0.011 | 0.021 | -0.54 | 67 | 0.59 |
| **Log-S:N** | **0.16** | **0.046** | **3.45** | 67 | **0.0010** |

#### *D: Full model excluding an outlier*

| | *Estimate* | *SE* | *t-value* | *df* | *P-value* |
|---|---|---|---|---|---|
| **Generations** | **0.0037** | **0.0018** | **2.08** | 66 | **0.042** |
| Gradient category (aridity) | 0.0097 | 0.24 | 0.04 | 66 | 0.97 |
| Gradient category (BioClim) | -0.42 | 0.30 | -1.37 | 66 | 0.17 |
| Gradient category (elevation) | -0.20 | 0.44 | -0.45 | 66 | 0.65 |
| Gradient category (temperature) | 0.11 | 0.25 | 0.42 | 66 | 0.68 |
| Mating system | 0.017 | 0.089 | 0.19 | 66 | 0.85 |
| Hemisphere | 0.12 | 0.12 | 1.07 | 66 | 0.29 |
| Gradient overlap | -0.73 | 0.44 | -1.67 | 66 | 0.10 |
| Trait number | -0.0042 | 0.021 | -0.20 | 66 | 0.84 |
| **Log-S:N** | **0.15** | **0.046** | **3.39** | 66 | **0.0012** |

**Table S5.** Tests of moderators on the relative magnitude of multivariate vector ($ln(||\mathbf{b_I}||/||\mathbf{b_N}||)$). The best model included no moderators in both analyses using all samples and without an outlier, indicating that our moderators explain little variation in $ln(||\mathbf{b_I}||/||\mathbf{b_N}||)$. The tables show the results of the full model with all samples (**A:** model fit: AICc = 156.9, $F_{10,67}$ = 1.2, P = 0.33, QE = 331.2), and without an outlier (**B**: AICc = 154.0, $F_{10,66}$ = 1.2, P = 0.32, QE = 328.4).

#### *A: Full model with all samples included*

| | *Estimate* | *SE* | *t-value* | *df* | *P-value* |
|---|---|---|---|---|---|
| Generations | 0.0058 | 0.0037 | 1.57 | 67 | 0.12 |
| Gradient category (aridity) | -0.75 | 0.56 | -1.35 | 67 | 0.18 |
| Gradient category (BioClim) | -0.42 | 0.58 | -0.72 | 67 | 0.48 |
| Gradient category (elevation) | -0.27 | 0.90 | -0.30 | 67 | 0.76 |
| Gradient category (temperature) | -0.35 | 0.58 | -0.61 | 67 | 0.55 |
| Mating system | 0.051 | 0.20 | 0.26 | 67 | 0.80 |
| Hemisphere | -0.12 | 0.18 | -0.67 | 67 | 0.51 |
| Gradient overlap | -1.0 | 0.68 | -1.47 | 67 | 0.15 |
| Trait number | 0.058 | 0.034 | 1.69 | 67 | 0.096 |
| Log-S:N | -0.03 | 0.073 | -0.42 | 67 | 0.68 |

#### *B: Full model excluding an outlier*

| | *Estimate* | *SE* | *t-value* | *df* | *P-value* |
|---|---|---|---|---|---|
| Generations | 0.0039 | 0.0041 | 0.96 | 66 | 0.34 |
| Gradient category (aridity) | -0.75 | 0.55 | -1.36 | 66 | 0.18 |
| Gradient category (BioClim) | -0.56 | 0.60 | -0.93 | 66 | 0.36 |
| Gradient category (elevation) | -0.47 | 0.92 | -0.50 | 66 | 0.62 |
| Gradient category (temperature) | -0.38 | 0.58 | -0.67 | 66 | 0.51 |
| Mating system | 0.071 | 0.20 | 0.36 | 66 | 0.72 |
| Hemisphere | -0.047 | 0.20 | -0.24 | 66 | 0.81 |
| Gradient overlap | -1.15 | 0.69 | -1.66 | 66 | 0.10 |
| Trait number | 0.064 | 0.035 | 1.83 | 66 | 0.072 |
| Log-S:N | -0.034 | 0.073 | -0.47 | 66 | 0.64 |

**Table S6.** Pooled effect sizes of multivariate parameters with different environmental gradients for estimating clines: *Authors' choice* uses the gradient chosen by each study, and *latitude* is based on absolute values of latitude where source populations originated from. *Temperature* and *precipitation* were extracted for each population to provide the same environmental metrics related to variability in temperature and seasonality (RCA1) and precipitation (RCA2) across all studies (Atwater et al., 2018). We first extracted 19 BIOCLIM variables for each population used in our study, and projected the varimax-rotated loadings, RCA1 and RCA2, from Atwater et al. (2018) on to the BIOCLIM values. The RCA1 and RCA2 loadings are appropriate for our study, as Atwater et al. (2018) used the climatic distribution of native and introduced pairs of 815 plant species, which includes our study species. 95% confidence intervals are shown in parentheses. $\mathbf{b_I}$ and $\mathbf{b_N}$ are vectors of introduced and native cline slope estimates. Cos($\theta$) represents the correlation of cline slopes between the ranges ($\theta$ being the angle between vectors), and $ln\|\boldsymbol{b_I}\|/\|\boldsymbol{b_N}\|$ shows the log ratio of the cline vector magnitudes. Asterisks indicate significance: * $P < 0.05$, ** $P < 0.01$, *** $P <0.001$.

| | $\cos(\theta)$ | $\ln(\|\boldsymbol{b_I}\|/\|\boldsymbol{b_N}\|)$ |
|---|---|---|
| Authors' choice | **0.30 [0.13, 0.46] ***** | **-0.30 [-0.58, -0.02] ***** |
| Latitude | **0.25 [0.067, 0.44] **** | **-0.22 [-0.47, 0.019]†** |
| Temperature | 0.14 [-0.06, 0.35] | **-0.30 [-0.60, -0.004] *** |
| Precipitation | -0.017 [-0.16, 0.13] | **-0.47 [-0.88, -0.06] *** |

**Table S7**. Effects of generations since introduction on multivariate parameters (see Table S5), with clines estimated using different environmental gradients. The slope of the relationship and 95% CI (in parentheses) are shown. Asterisks indicate significance: * $P < 0.05$, ** $P < 0.01$, *** $P < 0.001$.

| | $\cos(\theta)$ | $\ln(\lVert \boldsymbol{b_I} \rVert / \lVert \boldsymbol{b_N} \rVert)$ |
|---|---|---|
| Authors' choice | **0.0037 [0.0012, 0.0062]** ** | 0.0008 [-0.003, 0.0046] |
| Latitude | **0.0053 [0.0024, 0.0081]** *** | -0.0018 [-0.006, 0.0018] |
| Temperature | -0.0006[-0.0027, 0.0039] | -0.0056 [-0.011, -0.0006] * |
| Precipitation | -0.0013 [-0.041, -0.0014] | -0.0047 [-0.011, 0.002] |

**Table S8**. Univariate analysis of the pooled effect sizes of the relative magnitudes of clinal divergence for each trait category. $b_I$ and $b_N$ are cline slope estimates in the introduced and native ranges, respectively. Mean ln($b_I$/$b_N$) are shown, with 95% confidence intervals (CI) in brackets, and % magnitude of introduced range relative to native range in parentheses (% magnitude = exp(ln(x))*100). Effect sizes were estimated using traits with significant clines in at least one range and included positive values of $b_I$/$b_N$ only.

| | Mean ln($b_I$/$b_N$) | 95% CI | *t*-value | *P*-value |
|---|---|---|---|---|
| All traits | -0.22 (80%) | [-0.47, 0.02] | 1.8 | 0.08 |
| Defense | -0.01 (99%) | [-0.64, 0.61] | 0.04 | 0.97 |
| Reproductive | -0.04 (97%) | [-0.34, 0.27] | 0.23 | 0.81 |
| Morpho-physiology | -0.39 (68%) | [-0.82, 0.03] | 1.8 | 0.07 |
| Phenology | -0.27 (76%) | [-0.58, 0.04] | 1.7 | 0.09 |
| **Size** | **-0.33 (72%)** | **[-0.63, -0.03]** | **2.2** | **0.03** |

**Table S9.** Tests of moderators on the relative magnitude of the individual trait slopes ($\ln(|b_I/b_N|)$). The best model included no moderators, indicating that our moderators explain little variation in $\ln(|b_I/b_N|)$. The table shows the results of the full model.

| | *Estimate* | *SE* | *t-value* | *df* | *P-value* |
|---|---|---|---|---|---|
| Gene,rations (G) | 0.001 | 0.007 | 0.16 | 143 | 0.88 |
| Gradient category (aridity) | 0.25 | 0.37 | 0.68 | 143 | 0.50 |
| Gradient category (BioClim) | 0.48 | 0.74 | 0.65 | 143 | 0.52 |
| Gradient category (elevation) | 0.45 | 0.84 | 0.53 | 143 | 0.60 |
| Gradient category (temperature) | 0.31 | 0.38 | 0.83 | 143 | 0.41 |
| Mating system | 0.09 | 0.18 | 0.50 | 143 | 0.62 |
| Hemisphere | 0.03 | 0.23 | 0.13 | 143 | 0.90 |
| Gradient overlap | -0.03 | 0.91 | -0.04 | 143 | 0.97 |
| Trait category (defence) | -0.26 | 0.60 | -0.43 | 143 | 0.67 |
| Trait category (morpho-physio) | -0.34 | 0.48 | -0.71 | 143 | 0.48 |
| Trait category (phenology) | **-1.03** | **0.48** | **-2.15** | 143 | **0.03** |
| Trait category (reproduction) | -0.27 | 0.50 | -0.54 | 143 | 0.59 |
| Trait category (size) | -0.25 | 0.43 | -0.58 | 143 | 0.56 |
| Trait category (defence) x G | -0.01 | 0.01 | -0.71 | 143 | 0.48 |
| Trait category (morpho-physio) x G | 0.002 | 0.007 | 0.29 | 143 | 0.77 |
| Trait category (phenology) x G | -0.002 | 0.007 | -0.26 | 143 | 0.79 |
| Trait category (reproduction) x G | -0.005 | 0.007 | -0.63 | 143 | 0.53 |
| Trait category (size) x G | 0.001 | 0.007 | 0.16 | 143 | 0.88 |

**Table S10**. Relative change in *temperature* and *precipitation* (see Table 6) along the distance measures (i.e., latitude or altitude) between the pairs of introduced and native ranges. The pooled effect sizes of the relative slope ($\ln(b_I/b_N)$) and 95% CI were calculated using *rma.mv* function. The log ratio of the slopes overlapping with zero indicates that environmental gradient is similar between introduced and native ranges.

| | Mean $\ln(b_I/b_N)$ | 95% CI |
|---|---|---|
| Temperature | -0.06 | [-0.35, 0.22] |
| Precipitation | 0.26 | [-0.21, 0.72] |
